# Shotgun metagenomic and untargeted metabolomic analysis of gut microbiome associated with peripheral artery disease

**DOI:** 10.64898/2026.10.16.757428

**Authors:** Sarbjeet Niraula, Jae Woong Jang, Jon Jung, Spencer L. Stirewalt, Megan Alagna, James Du, Christopher Staley, Patrick Seed, Stefan J. Green, Karen J. Ho

**Affiliations:** Department of Surgery, Feinberg School of Medicine, Northwestern University, 676 North St. Clair Street, Suite 650, Chicago, IL, 60611, USA; Department of Surgery, University of Washington School of Medicine, 1959 NE Pacific Street, Box 356410, Seattle, WA 98112 USA; The University of Chicago Crown Family School of Social Work, Policy, and Practice, 969 East 60th Street, Chicago, IL 60637 USA; Department of Math, Statistics, and Computer Science, University of Illinois Chicago, 851 South Morgan Street, Chicago, IL 60607 USA; Department of Surgery, University of Minnesota, 420 Delaware Street SE Mayo Mail Code 195, Minneapolis, MN 55455; Department of Pediatrics, Division of Infectious Diseases, Ann & Robert H. Lurie Children’s Hospital of Chicago, Northwestern University Feinberg School of Medicine, Chicago, IL, USA; Genomics and Microbiome Core Facility, Rush University, Chicago, IL, USA

**Keywords:** Peripheral artery disease, humans, risk factors, cardiovascular diseases, dysbiosis, gastrointestinal microbiome, short chain fatty acids

## Abstract

**Background:** While the human gut microbiome is a potentially important and modifiable risk factor for cardiovascular diseases, the links between the gut microbiome and peripheral artery disease (PAD) and its clinical subgroups (claudication and chronic limb-threatening ischemia; CLTI) are not well understood.

**Methods:** We used shotgun metagenomics and untargeted metabolomics to analyze the fecal samples of people with PAD and age- and sex-matched non-PAD controls collected in a cross-sectional manner. We used functional prediction pathways, a network analysis, and weighted gene co-expression network analysis of metagenomic, metabolomics, and clinical data to better understand PAD-associated dysbiosis.

**Results:** Alpha and beta diversity were similar between the PAD (n=78) and non-PAD (n=20) groups. Differentially abundant species were dominated by those belonging to the Lachnospiraceae, Ruminococcaceae and Enterobacteriaceae families. Fecal butyrate levels were reduced between the non-PAD and PAD groups, but differences were not statistically significant. There was significantly reduced alpha diversity in the samples of the CLTI compared to claudication subgroup. Abundance of multiple butyrate producers were greater in the claudication group compared to CLTI. Network analysis revealed 5 meta-modules, each with specific associations to PAD presentation and metabolomic profiles, and species that are both top module hubs and differentially abundant between groups. Community interconnectedness appears to increase in claudication, followed by community disintegration in CLTI.

**Conclusions:** This metagenomic-metabolomic analysis of the gut microbiome of individuals with PAD, highlights the importance of studying functional capacity, diversity, and connectivity between species. Microbial modules may allow an integrative view of the microbial ecology relevant to presence and progression of PAD.

## Introduction

Lower extremity peripheral artery disease (PAD) is a form of cardiovascular disease that is a major cause of disability, mortality, and high healthcare costs. ^1-4^ Patients with PAD may experience exertional leg symptoms, functional impairment, progressive functional decline, and lower extremity rest pain, ulceration, and gangrene. ^5^ Global prevalence of PAD is projected to increase by 220% over 3 decades to reach 360 million by 2050. ^6^

While the high burden of PAD can be attributed to the increasing prevalence of traditional risk factors such as age, smoking, diabetes mellitus, hypertension, hyperlipidemia, and a sedentary lifestyle, only approximately 55% of PAD disease burden can be attributed to identified risk factors. ^7^ There are accumulating data that the human gut microbiome is a potentially important and modifiable risk factor for cardiovascular diseases, ^8^ but the links between the gut microbiome and PAD are not well understood. We previously found associations between plasma levels of specific microbiome-associated metabolites and the presence of PAD, ankle-brachial index and post-revascularization outcomes. ^9,10^ Trimethylamine N-oxide, a pro-atherogenic metabolite formed by gut microbes, also predicts long-term mortality in people with PAD. ^11^ In a gut microbiome GWAS dataset comprised of genome-wide genotypes and 16S rRNA amplicon sequencing data from 24 cohorts (total of 18,340 European individuals), the genera *Coprococcus 2*, *Ruminococcaceae UCG-004*, and *Ruminococcaceae UCG-010*, were identified as protective factors against PAD, while the Clostridiales cluster XI and the genera *Lachnoclostridium* and Lachnospiraceae UCG-001 were identified as risk factors for PAD. ^12^ However, there are substantial knowledge gaps in the involvement of the gut microbiome in PAD pathogenesis, clinical subtypes, outcomes.

To begin to address these gaps, we performed a metagenomics and untargeted metabolomic analysis of the gut microbiome in people with PAD and age- and sex-matched controls using a well-annotated clinical dataset. We characterized gut microbiome dysbiosis in PAD patients, identified species associated with progressive clinical symptoms of PAD, analyzed functional prediction pathways to infer microbial processes associated with dysbiosis in PAD, and performed a network analysis of metagenomic sequencing, metabolomics, and clinical data to better understand these interactions.

## Methods

### 1. Study participants

This is a retrospective study of study participants aged ≥ 40 years with documented PAD and non-PAD controls who were recruited from a single vascular surgery practice at a single academic medical center between 2021 and 2023. Exclusion criteria included inability to provide informed consent or follow the study protocol, antibiotic use within the past 3 months, or bowel surgery or bowel preparation within the past 3 months. After obtaining informed consent, instructions about home fecal sample collection and a collection kit were provided. Demographic and clinical information were obtained from the electronic medical record or a health questionnaire. This study was conducted in accordance with the Declaration of Helsinki and approved by the Northwestern University Institutional Review Board (Protocols #STU00213201 and #STU00216722).

### 2. Sample collection and processing

All participants were instructed to freeze fecal samples immediately after collection and transport them back to the study team in an insulated bag with an ice pack. Upon return to the study team, samples were stored immediately at -80°C until batch DNA extraction. DNA was extracted using the PowerSoilPro DNA isolation kit (Qiagen, Germantown, USA). Shotgun metagenome library preparation was performed using an Illumina DNA prep kit, according to the manufacturer’s instructions. Five cycles of PCR were performed for all samples. Libraries were pooled and sequenced on an Illumina NovaSeq X instrument, employing a 10B flow cell lane and 2x150 base sequencing. Library preparation was performed at the Genomics and Microbiome Core Facility at Rush University (RRID:SCR_026856), and sequencing was performed at the DNA Services Facility, Roy J. Carver Biotechnology Center, University of Illinois Urbana-Champaign.

### 3. Bioinformatics analysis

Raw paired-end sequences were quality filtered using the KneadData (v0.12.0) ^13^ pipeline, which included Trimmomatic (v0.39) ^14^ for adapter removal and Bowtie2 (v2.5.2) ^15^ for filtering human read contaminants from human reference database (hg37dec_v0.1). Taxonomic annotation was performed using Kraken2^16^ against the Unified Human Gastrointestinal Genome (UHGG) database (v2.0.2). ^17^ Confidence and minimum-hit-group threshold for Kraken2 was setup as 0.3 and 3, respectively. Bracken (v2.9) ^18^ was used to estimate the abundance of species from the Kraken2 report file with minimum read number 10 and the default read length of at least 100. Quality filtered paired and unpaired reads were concatenated, followed by functional prediction using the Humann3.9^19^ pipeline with default settings. Gene prediction was done using the UniRef90 (v201901b_full) database) ^20^ and CHOCOPhlAn (v201901_v31). ^21^ We also selected MetaPhlAn database version (mpa_vJun23_CHOCOPhlAnSGB_202403) ^22^ for annotation. MetaCyc pathway abundances^23^ were transformed to copies per million (CPM) for further analysis.

### 4. Untargeted metabolomics

Fecal samples were lyophilized in the Northwestern University Analytical BioNanoTechnology Equipment Core (RRID: SCR_023706) and freeze-dried samples were weighed prior to untargeted metabolite quantification using nuclear magnetic resonance (NMR). The metabolomics analysis was performed at the Cincinnati Children’s Hospital NMR based Metabolomics Core Facility (RRID: SCR_022636) according to the previously published protocol. ^24^ Briefly, dried fecal material (100 mg ± 5%) was homogenized in cold phosphate buffered saline and the supernatant was filtered using pre-washed 3 kDa spin filters (NANOSEP 3K, Pall Life Sciences, Portsmouth, UK). The NMR sample was prepared with 300 µL of fecal filtrate with NMR buffer containing an internal standard, trimethylsilylpropanoic acid (TMSP). The final sample volume was 600 µL and final TMSP concentration in each sample was 0.5 mM. NMR data collection and processing were performed with a Bruker Avance III HD 600 MHz spectrometer using Topspin 3.6 software (Bruker Analytik, Rheinstetten, Germany) as previously described. ^25^ A total of 41 metabolites were assigned based on the chemical shifts on 1D ^1^H-NOESY, 2D-TOCSY and -HSQC NMR experiments with reference spectra found in databases, Human Metabolome Database (HMDB), ^26^ and Chenomx® NMR Suite profiling software (Chenomx Inc. version 8.1). The metabolites were quantified by Chenomx software based on the internal standard TMSP and were normalized to sample weights prior to statistical analysis.

### 5. Network analysis

SparCC ^27^ correlation networks of species, with at least 10% prevalence within each group (non-PAD, PAD, and the PAD subgroups) were constructed with FastSpar^28^ using default parameters. P values were calculated using 1000 permutations.^29^ We selected edges with significant (p ≤ 0.005) and strong (correlation threshold ≥ 0.3) relationships. ^27^ Nodes (species) common in all four networks were selected for further analysis. The unsigned topological overlap matrix was created from the network with zero negative associations. Modules were created using average clustering with minimum module size of 25 taxa and deep split 2 in the weighted gene co-expression network analysis (WGCNA) package. ^30^ Module eigengenes (ME), the first principal component of each module, were Spearman correlated with sample traits in the WGCNA package. ^30^ Metabolite concentrations were log-transformed and mean-centered. PAD severity was converted to numerical values before correlation. ME were correlated and ME distances were clustered using hierarchical clustering. Module abundances were calculated by summing the abundances of species in each module.

To evaluate whether PAD network modules were also present in the non-PAD network, we used the modulePreservation function within the WGCNA package with 1000 permutations. The Zsummary score >10 represents high preservation, 2 < Zsummary < 10 represents moderate preservation and Zsummary <2 is no preservation. ^31^ Similarly, lower median rank represents better relative preservation, which is independent of module size. The weighted hub score of taxa in the network was calculated using igraph package^32^ and module hubs were defined as top 25 percentile of taxa with highest intramodular connectivity Z score (kWithin_z) within each module. Network visualization was done using Cytoscape (v 3.20.2) ^33^ and the module circos plot was created using circlize R package (v 0.4.16). ^34^ Abundance data and statistical analyses were performed in R.

### 6. Statistical analysis

Differential abundance analysis was performed using the ANCOM-BC2 package^35^ in R. Species with at least 10 reads in at least 20% of the samples within each group were considered for differential abundance analysis. We adjusted for age, sex, race, body mass index (BMI), smoking, and history of coronary artery disease and diabetes mellitus in all models. We considered differential abundance for species that have Benjamini-Hochberg (BH)-adjusted p values ≤ 0.05 and passed the sensitivity analysis test for pseudo count addition.

Alpha diversity was calculated using observed richness, Shannon, Pielou evenness and Simpson indices. Observed richness was calculated from raw count data and adjusted for sequencing depth using the mstat_alpha_diversity_adjusted function in the MicrobiomeStat (v1.2.1) package. ^36^ The other 3 alpha diversity indices were calculated using relative abundance data. The Wilcoxon rank sum test was used to compare alpha diversities between groups and BH-adjusted for multiple comparisons.

Beta diversity was estimated using Robust Aitchison principal component analysis (rPCA) using the Gemelli R package. ^37^ Differences between groups were based on the non-parametric permutational multivariate ANOVA (PREMANOVA) test with the vegan^38^ R package using 9999 permutations.

Differential abundance of MetaCyc functional pathways were compared using a generalized linear model on centered log ratio-transformed abundance data. P values are reported without adjustment.

In the network analysis, genus enrichment in each module was performed using the Fisher’s exact test and p values were BH-adjusted. Network analysis was done using igraph^32^ and WGCNA packages in R.

### 7. Independent data access and analysis

We attest that one author (KH) had full access to all the data in the study and takes responsibility for its integrity and the data analysis.

## Results

### 1. Study cohort

There were 98 participants, of whom 78 had PAD and 20 were non-PAD controls. The mean age of the PAD group is 70.7 years (standard deviation [SD] 9.4; range 49-90), which is not significantly different from the non-PAD controls (mean age 65.0 years; SD 12.7, range 40-82). There were more females in the non-PAD group (n=13, 65%) compared to the PAD group (n=35, 44.9%), but this was also not statistically significant. As expected, 35 (44.9%) participants in the PAD group have diabetes, 64 (82.1%) have hypertension, 42 (53.9%) have coronary artery disease, and 65 (83.3%) are current or former smokers. Of those with PAD, 38 (48.7%) had symptoms of claudication and 39 (51.3%) had chronic limb threatening ischemia (CLTI). Of those with CLTI, 18 (45%) had rest pain, and 21 (30.5%) had non healing ulcers or gangrene. The demographic and clinical characteristics of both groups are shown in **Table 1**.

**Table 1.**
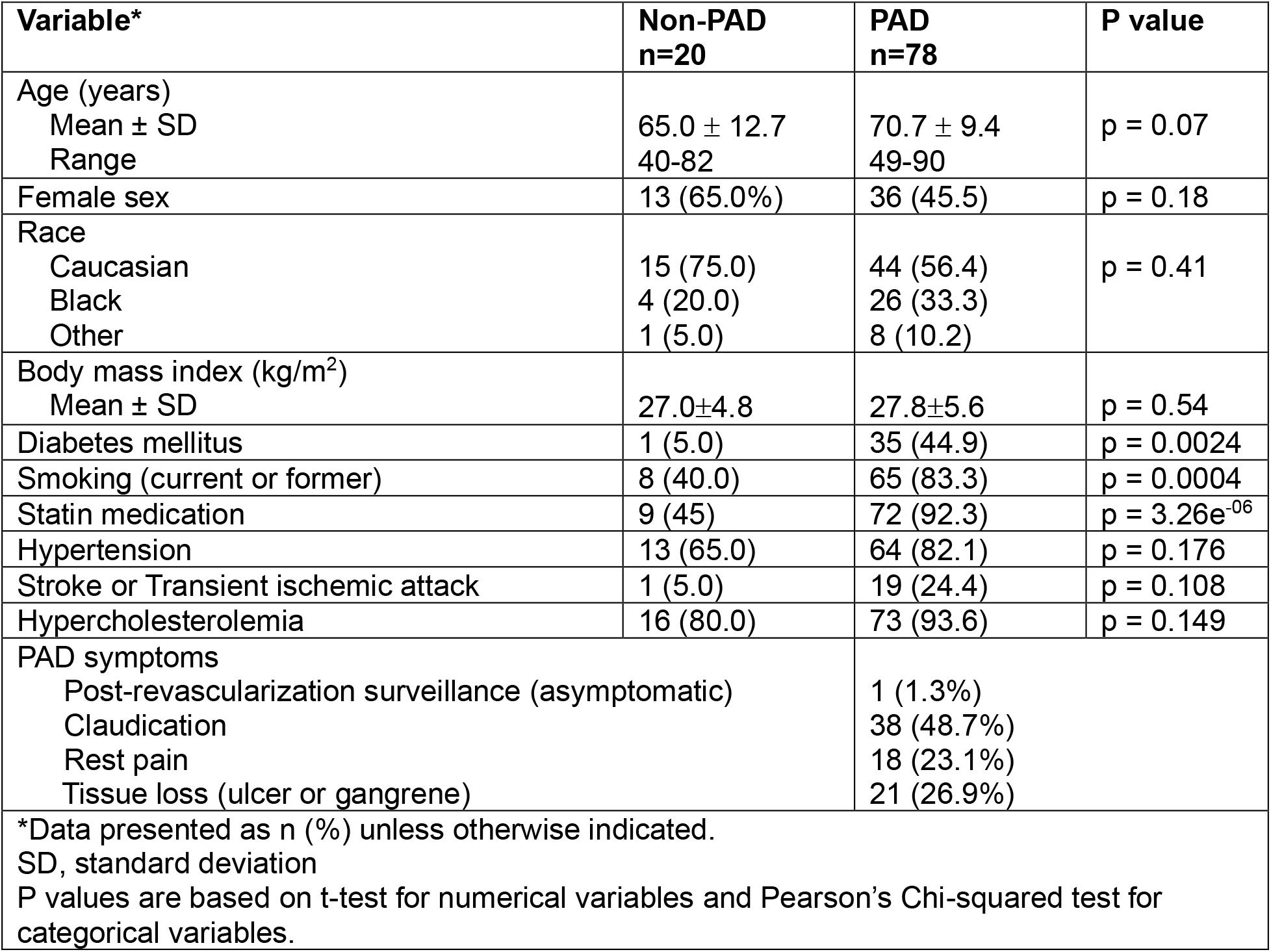
Demographic and clinical variables.

| Variable* | Non-PAD<br>n=20 | PAD<br>n=78 | P value |
| --- | --- | --- | --- |
| Age (years)<br>Mean ± SD<br>Range | 65.0 ± 12.7<br>40-82 | 70.7 ± 9.4<br>49-90 | p = 0.07 |
| Female sex | 13 (65.0%) | 36 (45.5) | p = 0.18 |
| Race<br>Caucasian<br>Black<br>Other | 15 (75.0)<br>4 (20.0)<br>1 (5.0) | 44 (56.4)<br>26 (33.3)<br>8 (10.2) | p = 0.41 |
| Body mass index (kg/m<sup>2</sup>)<br>Mean ± SD | 27.0±4.8 | 27.8±5.6 | p = 0.54 |
| Diabetes mellitus | 1 (5.0) | 35 (44.9) | p = 0.0024 |
| Smoking (current or former) | 8 (40.0) | 65 (83.3) | p = 0.0004 |
| Statin medication | 9 (45) | 72 (92.3) | p = 3.26e<sup>-06</sup> |
| Hypertension | 13 (65.0) | 64 (82.1) | p = 0.176 |
| Stroke or Transient ischemic attack | 1 (5.0) | 19 (24.4) | p = 0.108 |
| Hypercholesterolemia | 16 (80.0) | 73 (93.6) | p = 0.149 |
| PAD symptoms<br>Post-revascularization surveillance (asymptomatic)<br>Claudication<br>Rest pain<br>Tissue loss (ulcer or gangrene) |  | 1 (1.3%)<br>38 (48.7%)<br>18 (23.1%)<br>21 (26.9%) |  |
| *Data presented as n (%) unless otherwise indicated.<br>SD, standard deviation<br>P values are based on t-test for numerical variables and Pearson's Chi-squared test for categorical variables. |  |  |  |

### 2. Alpha and beta diversity

After quality filtering and host contaminant removal using the KneadData pipeline, the remaining high-quality reads were aligned using Bowtie2, ^15^ which revealed that the proportions of human host contaminant reads were 2.12 ± 0.81% and 1.83 ± 0.76% in the non-PAD and PAD groups, respectively (**Supplemental Table 1**). Alpha diversity indices were all similar between the non-PAD and PAD groups overall (**Figure 1A**). Among males, alpha diversity in the PAD group was higher than in the non-PAD group. Among females, alpha diversity was higher in the non-PAD group. However, none of these differences were statistically significant (**Supplemental Figure 1A**).

**Figure 1.**
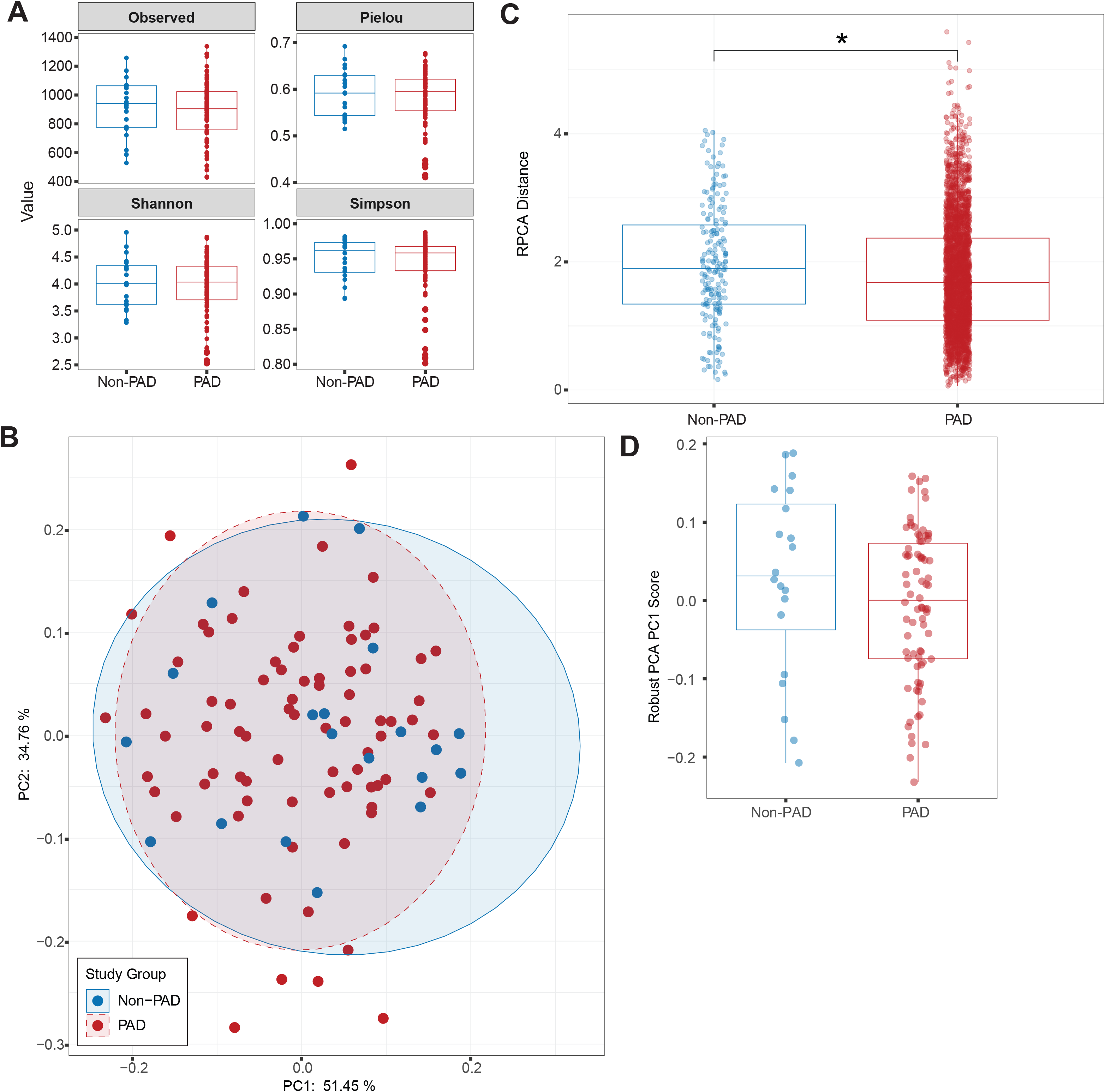
Alpha and beta diversity. (A) Alpha diversity metrics, including observed richness, Pielou evenness, Shannon, and Simpson indices, were compared between the non-PAD and PAD groups. (B) Robust Aitchison principal coordinate analysis (RPCA) plot of samples from the non-PAD and PAD groups. Percentage values represented on the main axes indicate explained variance associated with each principal component. Shaded ellipses indicate 95% confidence regions around group centroids. Statistical differences between groups were assessed using PERMANOVA (R^2^=0.02, F= 2.37, p=0.1, permutations=9999). (C) RPCA distances between samples within the non-PAD and PAD groups, indicating that the non-PAD group is more heterogeneous. *, p = 0.006; Wilcoxon rank-sum test. (D) Comparison of PC1 sample scores from (B) between the non-PAD and PAD groups, indicating that there is similar variance of non-PAD and PAD samples in PC1 (Wilcoxon rank-sum test).

Beta diversity was similar between the non-PAD and PAD groups (**Figure 1B**), but the median distance within non-PAD samples was higher than within PAD samples (p = 0.006; **Figure 1C**), suggesting that the intra-group microbial community is more homogeneous among PAD participants compared to the non-PAD controls. Non-PAD samples had higher coordinate scores along the first principal component (PC1) (**Figure 1D**), which contributes to 51.45% of the variance in taxonomic composition (**Figure 1B**). However, the observed trend was not statistically significant due to high variability and substantial overlap between two groups.

### 3. Taxonomic characterization and differential abundance

Overall, 92.31 ± 2.67% of quality filtered reads were classified against the UHGG2 database, resulting in the identification of 3,531 species with at least 10 classified reads (**Supplemental Table 1**). We identified a total of 20 phyla in the samples. As shown in **Figure 2A**, 12 families comprised approximately 90% of the total abundance in both groups. Lachnospiraceae was the most abundant family (non-PAD 52.48 ± 15.24%; PAD 53.57 ± 18.18%), followed by Bacteroidaceae (non-PAD 9.09 ± 11.79%; PAD 6.83 ± 4.58%) and Ruminococcaceae (non-PAD 6.21 ± 3.25%; PAD 6.1 ± 7.53%). There were no significant differences in the abundances of families between the PAD and non-PAD groups.

**Figure 2.**
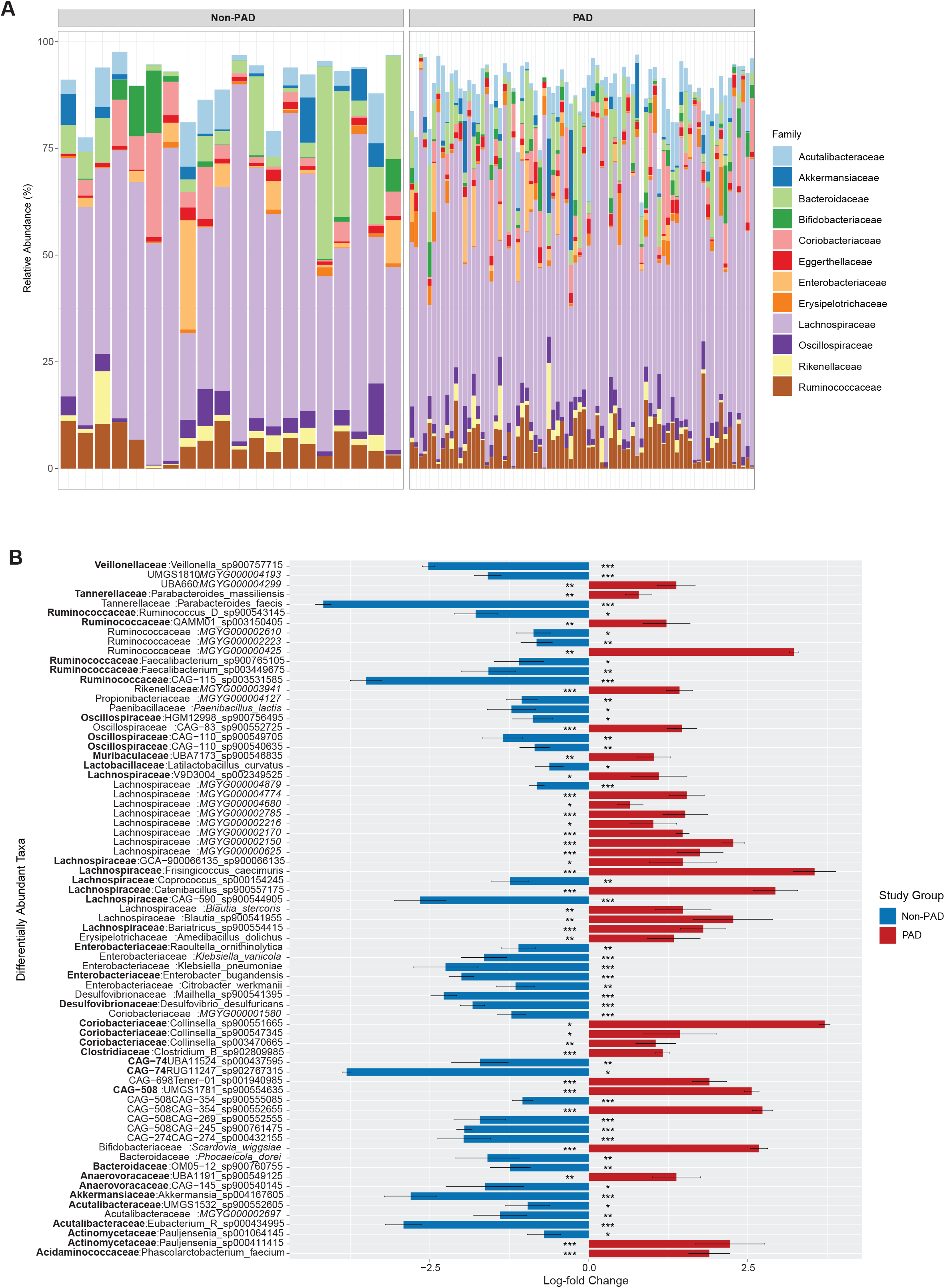
Taxonomy and differential relative abundance. (A) Taxonomic bar plot showing relative abundance of families in the non-PAD and PAD groups. Each bar represents an individual sample. (B) Differential abundance of species in the non-PAD and PAD groups. Taxa are grouped according to family. Taxa were considered differentially abundant if FDR-adjusted p values were significant (p_adj_ < 0.05) and they passed the sensitivity analysis test. *, p < 0.05. **, p < 0.01. ***, p < 0.001. Error bars represent standard deviation. FDR, false discovery rate.

There were multiple differentially abundant bacterial species between the non-PAD and PAD groups. Differentially abundant species were dominated by those belonging to the Lachnospiraceae, Ruminococcaceae and Enterobacteriaceae families (**Figure 2B**). All 5 differentially abundant species from Enterobacteriaceae, *Akkermansia_sp004167605, Latilactobacillus. Ruminococcus D sp900543145,* and *Parabacteroides faecis* were less abundant in the PAD group. On the other hand, *Colinsella* spp., which have been associated with inflammatory states, ^39,40^ diabetes, ^41^ and atherosclerosis, ^42^ was increased in the PAD group (**Figure 2B**).

### 4. Untargeted metabolomics

A total of 41 metabolites from 52 fecal samples (9 non-PAD and 43 PAD) were quantified, as shown in (**Supplemental Figure 2A**). On bivariate analysis, butyrate levels were reduced in the PAD compared to the non-PAD group (**Supplemental Figure 2A**), but this and other differences in metabolite levels between the non-PAD and PAD groups did not reach statistical significance. There were also no significant differences in short chain fatty acid levels between the claudication (n=24) and CLTI subgroups (n=19) or between the CLTI subgroups (n=9 rest pain, n=9 tissue loss and n=1 post revascular surveillance) (**Supplemental Figure 2B**).

### 5. Alpha and beta diversity within the PAD clinical subgroups

We next examined dysbiosis associated with the clinical subgroups of PAD, claudication and CLTI. There was reduced richness (p^adj^ = 0.011), evenness (p^adj^ = 0.031) and Shannon diversity (p^adj^ = 0.022) in the CLTI subgroup compared to the claudication subgroup (**Figure 3A**), suggesting an inverse association between microbial diversity with PAD severity. Male patients with CLTI had reduced alpha diversity (observed richness, Pielou evenness and Shannon diversity) compared to male patients with claudication, but there were no differences between female patients with claudication and CLTI (**Supplemental Figure 1B**), indicating possible sex-specific effects of PAD severity on diversity. There were also qualitative but non-significant declines in median alpha diversity estimates as severity of the presenting symptom of PAD increased (**Figure 3B**).

**Figure 3.**
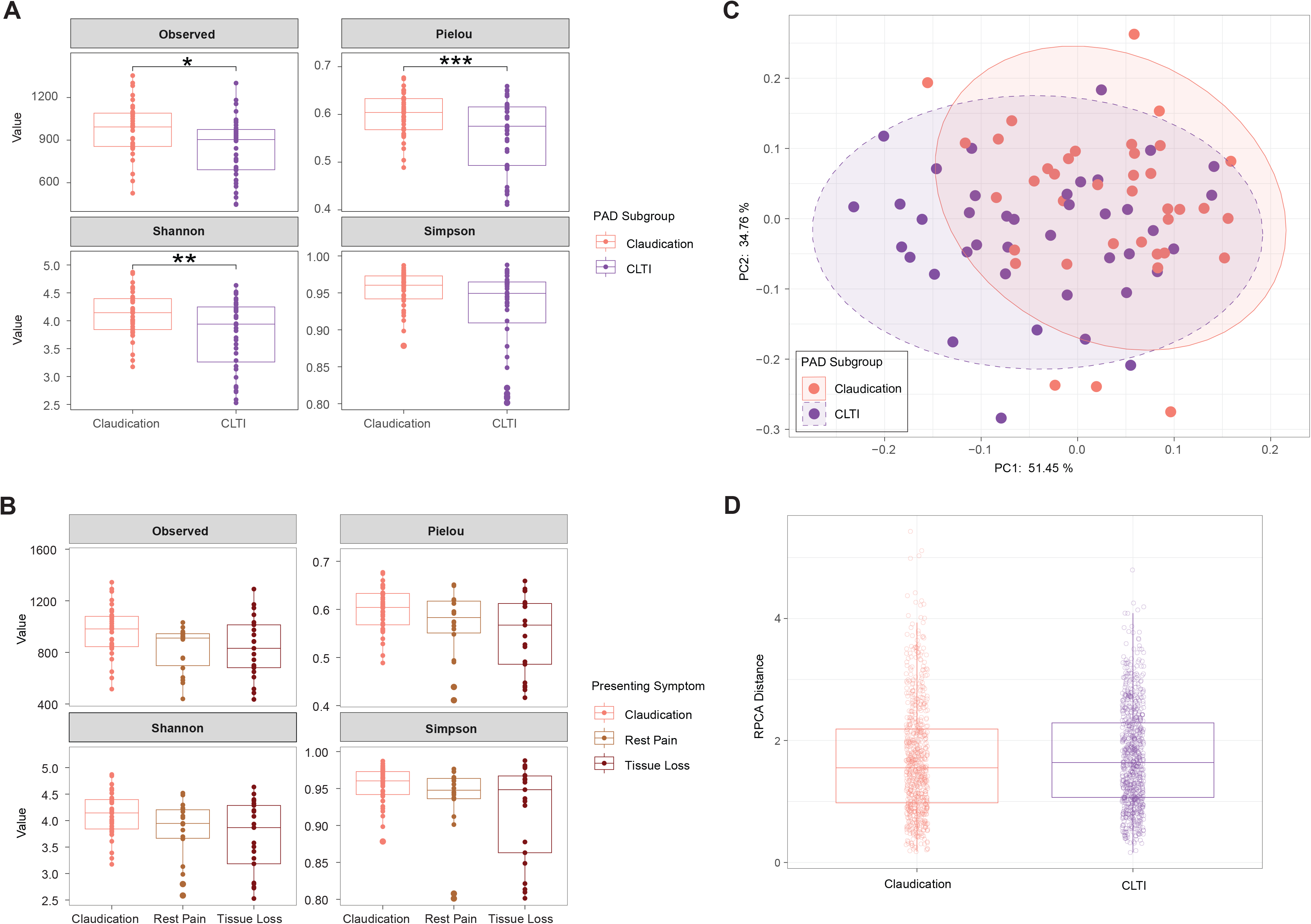
Alpha and beta diversity in subclinical groups of PAD. (A) Alpha diversity indices in claudication and chronic limb-threatening ischemia (CLTI) subgroups were compared. *, p = 0.01, **, p = 0.03, ***, p = 0.02,, Wilcoxon rank-sum test. (B) There were no significant differences in alpha diversity indices (Wilcoxon rank-sum test) between claudicants and those with rest pain or tissue loss. (C) RPCA plot between claudication and CLTI subgroups. Statistical differences between groups were assessed using PERMANOVA (R^2^=0.07, F=6.1, p=0.002, permutations=9999). (D) There were no significant difference in RPCA distances between samples in the claudication and CLTI subgroups (Wilcoxon rank-sum test). Alt-text: Graphical representation of alpha and beta diversity comparisons between the subgroups of claudication and CLTI, including statistical analyses.

As shown in **Figure 3C**, beta diversity (microbial composition) was overall different between the claudication and CLTI subgroups but homogeneity of microbial composition within the claudication and CLTI subgroups was similar (**Figure 3D**), suggesting a consistent shift in the gut microbial profiles in patients based on clinical presentation of PAD. The sample score in PC1 of the claudication subgroup was significantly higher than that of the CLTI subgroup (p^adj^ = 0.003) (**Figure 4A**) and specifically the CLTI presentation of tissue loss (p^adj^ = 0.002) (**Figure 4B**), suggesting that along PC1 (**Figure 1B** and **1D**), there is a compositional gradient associated with PAD severity rather than overall contrast between non-PAD and PAD. To identify the species driving variance in PC1, we created a list of the top 100 species loadings in PC1 (**Figure 4C**).

**Figure 4.**
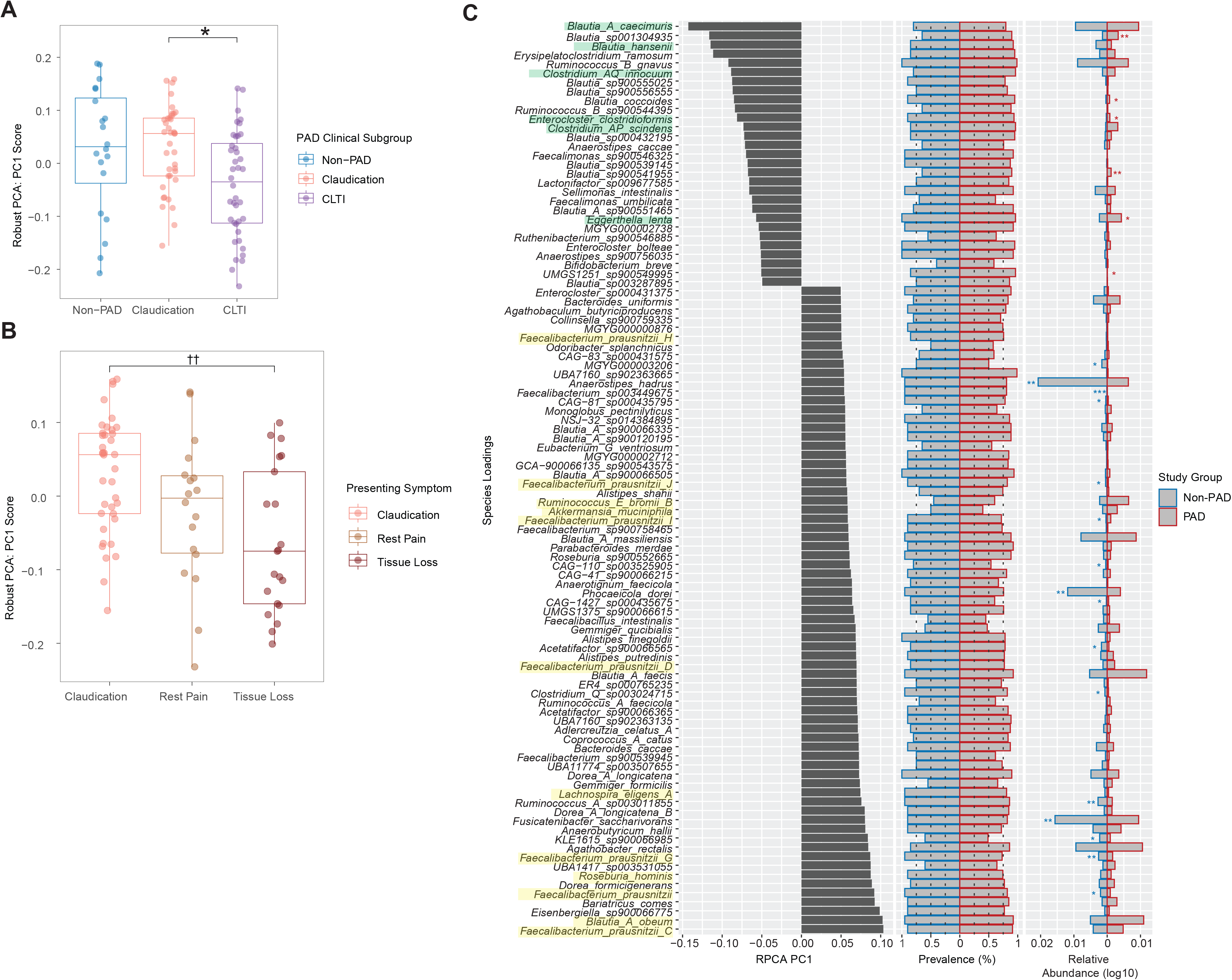
Principal component 1 sample scores in the non-PAD, claudication, and CLTI groups and taxa driving group separation. (A) Comparison of PC1 scores between the non-PAD, claudication and CLTI groups in the first principal component axes. *, p = 0.003, Wilcoxon rank-sum test. (B) Comparison of PC1 scores between the claudication, rest pain and tissue loss. ꝉꝉ, p = 0.002, Wilcoxon rank-sum test. (C) Contribution of the top 100 species to PC1 and their prevalence and abundance in the non-PAD and PAD groups. Species are ranked in descending order of absolute scores. Prevalence of each species is similar in both groups. Log-transformed relative abundance of each species between the groups was compared using the Wilcoxon rank-sum test. Species highlighted in yellow are known to be high butyrate producers that are positively associated with PC1. *, p < 0.05. **, p < 0.01. ***, p < 0.001.

Most of these species have low abundance in both PAD and non-PAD groups, but they show similar prevalence across comparison groups (**Figure 4C**). Among them, known butyrate producers^43^ such as *Faecalibacterium prausnitzii, Blautia A obeum, Blautia A faecis, Roseburia hominis, Lachnospira eligens A,* resistant starch degrader *Ruminococcus bromii* which leads to enhanced butyrate production, ^44-46^ and propionate and acetate producer *Akkermansia muciniphila*^47^ were positively associated with PC1 (indicated with yellow highlights in **Figure 4C**), suggesting that the relative abundance of these species correlates with non-PAD status. *Clostridium AP scindens, Enterocloster clostridioformis, Clostridium AQ innocuum and Eggerthella lenta,* were negatively associated with PC1 (indicated with green highlights in **Figure 4C**), suggesting that these species, which are known to be involved in secondary bile acid production, bile acid oxidation, and impaired intestinal barrier function, ^48-50^ may be associated with PAD. Similarly, we found a negative contribution of known acetate and succinate producers such as *Blautia hansenii* and *Blautia A caecimuris* (**Figure 4C**) on PC1, ^6^ suggesting a relative enrichment of these species in PAD.

### 6. Differential abundance in the PAD clinical subgroups

There were notable differences in bacterial abundance between claudication and CLTI, as shown in **Figure 5A**. Abundance of multiple butyrate producers such as *Roseburia hominis, Faecalibacterium prausnitzii C*, and *Eubacterium* spp. were more abundant in the claudication group compared to CLTI (**Figure 5A**). *Methanobrevibacter smithii*, which converts hydrogen from bacterial fermentation to methane^51^ and may participate in the regulation of butyrogenesis in the human colon, was also more abundant in the claudication group. ^52^ *Alistipes* spp., which were found to be less abundant in people with atherosclerosis compared to healthy controls in a different study, ^53^ was more abundant in the less clinically advanced PAD group in the current study. *Collinsella aerofasciens*, an actinomycete that is associated with chronic inflammatory conditions such as Crohn’s disease, ^54^ nonalcoholic fatty liver disease, ^55^ and symptomatic atherosclerosis, ^42^ was also more abundant in the claudication group, raising the hypothesis that it could be a biomarker of early-stage PAD. Similarly, *Desulfovibrio fairfieldensis*, a Deltaproteobacteria that produces hydrogen sulfide gas and has been found to be enriched in patients with inflammatory bowel disease, diabetes-associated coronary artery disease, and colon cancer, ^56^ was found to be more abundant in the earlier stage of PAD.

**Figure 5.**
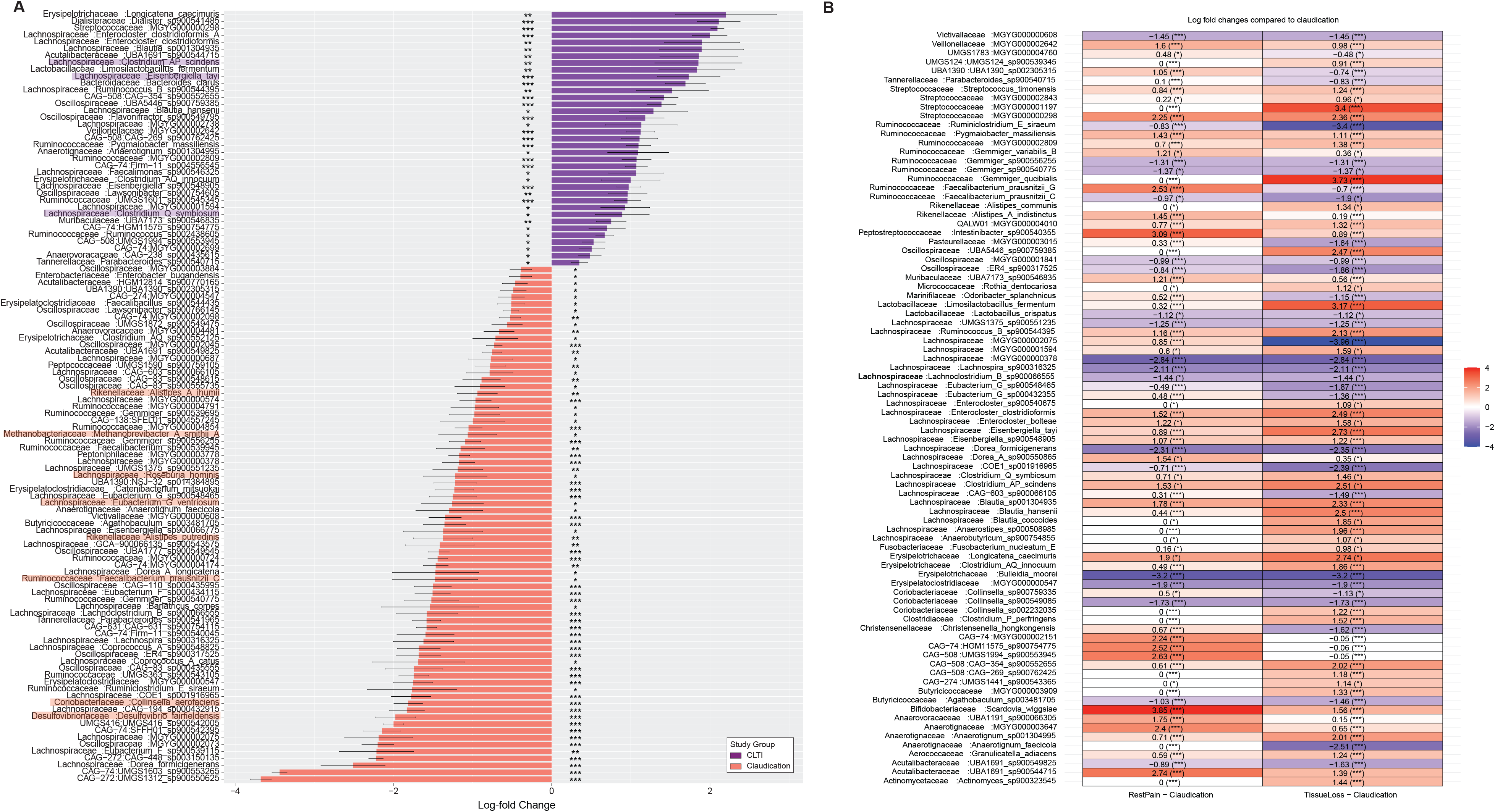
Differential relative abundance of species in the claudication and CLTI subgroups. (A) Differential abundance of species in the CLTI subgroup compared to claudication identified using ANCOM-BC2. We included species present in at least 20% of the samples. P values were Benjamini-Hochberg (BH)-adjusted. Species are presented in the descending order of the log fold change values. (B) Trend test of species in samples based on CLTI symptoms (rest pain or tissue loss) compared to claudication. Log fold changes are presented. P values are BH-adjusted. Error bars represent standard deviation.

There were fewer species that were differentially more abundant in CLTI compared to claudication (**Figure 5A**). These included *Clostridium scindens*, which produces secondary bile acids; ^50^ *Clostridium symbiosum*, which is associated with amino acid metabolism in colonocytes, ^57^ colon cancer, ^58^ coronary artery disease, ^59^ and irritable bowel syndrome; ^60^ and *Eisenbergiella tayi,* which is associated with intestinal inflammation and multiple sclerosis. ^61^ These bacteria showed a progressively higher abundance trend with increasing PAD severity (rest pain to tissue loss) relative to claudication (**Figure 5B**). In contrast, butyrate producers such as *Ruminoclostridium E siraeum* (p^adj^ < 0.001) and *F. prausnitzii G* (p^adj^ < 0.05) showed a progressively reduced abundance trend whereas *F. prausnitzii C* showed non-linear abundance change (**Figure 5B**).

### 7. PAD network analysis

In order to better understand the community interactions and microbiome associations in PAD, 4 networks consisting of common 1644 species (nodes) were constructed that have 23,108, 13,585, 16,905 and 14,298 interactions (edges) in non-PAD, PAD, claudication and CLTI samples, respectively. Overall, the node degree is higher in the claudication network compared to non-PAD and CLTI, whereas the clustering coefficient is reduced with increased PAD severity (**Figure 6A**), which suggest a significant rewiring of gut community structure at different stages of the disease.

**Figure 6.**
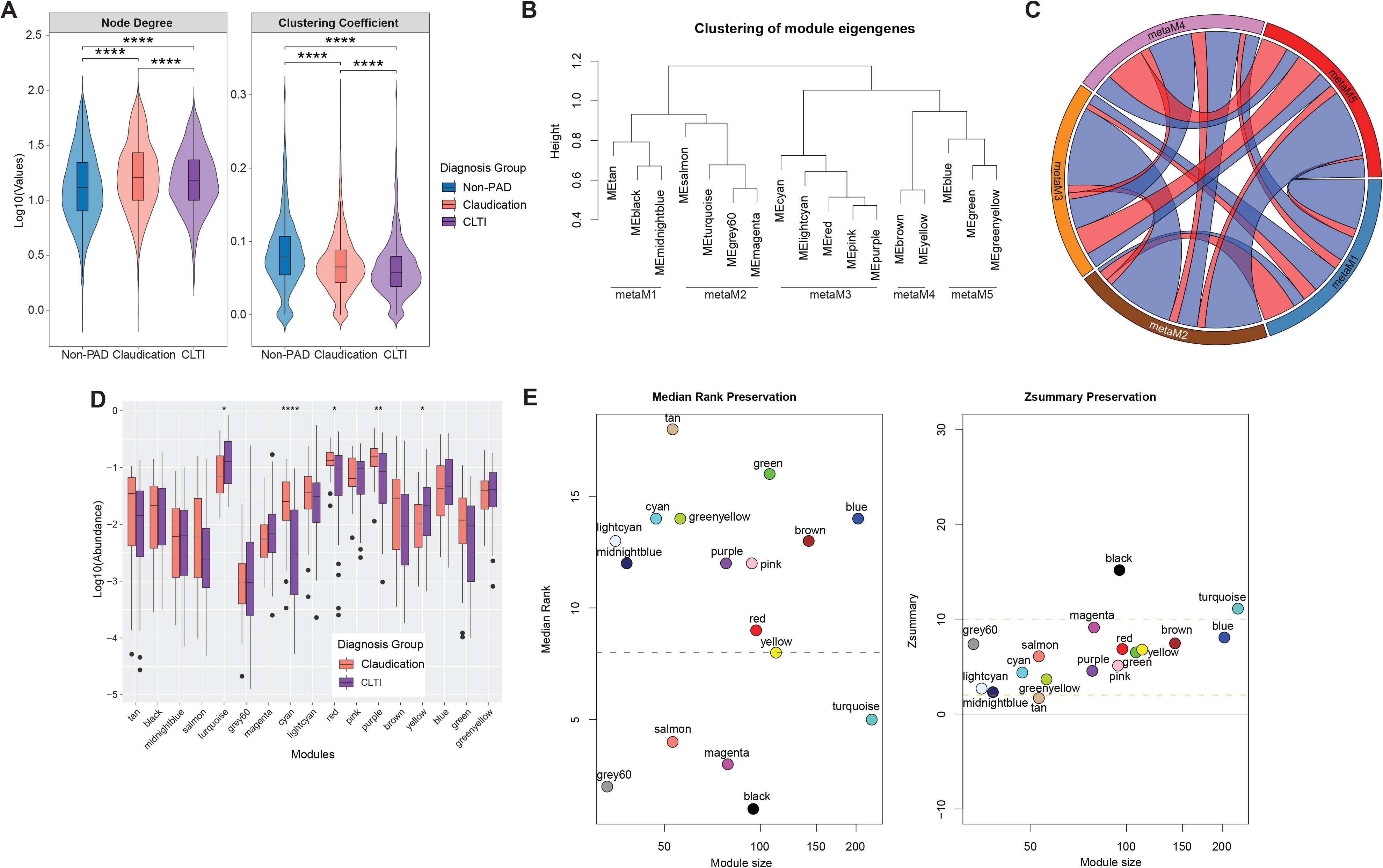
Species co-occurrence network analysis in non-PAD and PAD groups. (A) Properties of networks associated with the non-PAD, claudication, and CLTI groups. (B) Average linkage clustering dendrogram of 17 PAD network module clusters, which were further grouped into 5 meta-modules based on the clustering height of 0.9. ME, module eigengenes. (C) Circose plot showing the average number of positively (red) and negatively (blue) correlated edges (connections) between PAD network meta-modules. Ribbon width indicates the number of edges (at least 10) and opacity indicates average correlation strength. (D) Box and whiskers plot of the cumulative abundance of species within each module in the claudication and CLTI groups. *, p<0.05. **, p<0.01. ****, p<0.001, Wilcoxon rank-sum test. (E) Preservation of PAD network module connections within the non-PAD network based on median rank and Zsummary preservation statistics. Blue dashed line in left panel represents a median rank threshold of 8. Red and blue dashed lines in right sub-panel represent lower (Z=2) and upper (Z=10) threshold values for moderate z-statistics.

To achieve a higher-order organization of PAD network, species were clustered into eighteen modules, each assigned a different color. The grey module, which consists of species of undefined module membership, was excluded from further analysis (**Figure 6B**). Seventeen valid modules were further grouped into 5 meta-modules based on the distance between ME (**Figure 6B**). In our ME-based hierarchical clustering tree (**Figure 6B**), closely assigned modules and meta-modules share higher proportions of overall positive correlations between species and vice versa **(Figure 6C and Supplemental Figure 3A and 3B**), validating that the ME of each module is representative of the overall co-abundance pattern of the respective module members.

We next evaluated the structural organization and biological integrity of the modules. Module membership (kME) was strongly correlated (p < 0.05) with intramodular connectivity (kWithin), suggesting that the modules contain well defined communities with structural and functional coherence (data not shown). The kWithin of all modules, except those in meta-module 1 and salmon and grey60 from meta-module 2, have high correlation (p < 0.05) with intermodular connectivity (kOut), indicating that the module hubs that have more connections within the module also act as connectors between communities (data not shown). Conversely, modules lacking correlation between kWithin and kOut, such as tan, black, salmon and grey60, suggest a decoupling of local and global network roles.

Several modules are associated with PAD severity and fecal metabolites. Meta-module 3 members such as the “cyan,” “red”, “pink” and “purple” modules correlate with claudication (**Supplemental Figure 3C**; p < 0.05, Spearman correlation). The majority of these modules are also positively correlated with fecal butyrate, nicotinate, thiamine, uracil, formate, UDP-glucose and glutamate levels. The pink module also correlates positively with propionate. Butyrate and nicotinate are metabolites known for their protective role in atherosclerotic cardiovascular diseases. ^62-63^ The turquoise and magenta modules in meta-module 2 and blue module in meta-module 5 correlate with CLTI (**Supplemental Figure 3C**; p < 0.05), and they have negative connections with modules in meta-module 3 (**Supplemental Figure 3A and 3B**), suggesting that these modules are directly associated with PAD severity. The magenta module is negatively correlated with butyrate, isovalerate and inosine levels, whereas the turquoise module is positively correlated with acetate and propionate and 13 out of 14 of the measured amino acids. The green and greenyellow modules are negatively correlated with acetate, butyrate and propionate (**Supplemental Figure 3C**; p < 0.05). Cumulative relative abundances of species in the cyan, red and purple modules are lower in the CLTI compared to the claudication group, whereas those in the turquoise and yellow modules are comparatively more abundant in the CLTI group. (**Figure 6D**).

Modules associated with PAD and associated sample traits exhibited structured connectivity patterns, reflecting an altered state of community organization in PAD. Therefore, we compared the structural organization of PAD modules within non-PAD network. Module preservation statistics revealed that the modules within meta-module 2 are highly preserved in the non-PAD network (**Figure 6E**), suggesting that the connectivity pattern among members within those modules remains similar in the non-PAD group. Modules such as pink and red that are correlated inversely with PAD severity and positively with butyrate levels are relatively less preserved (Zsummary < 10 and median rank >8) in the non-PAD network than those with the opposite trend, such as magenta (**Figure 6E**). This suggests that interactions among species beneficial to health reorganize differently in the PAD compared to the non-PAD groups mainly by increasing internal cohesiveness (gain of density) and losing global network connections, as shown by lower Zdensity and higher Zconnectivity (**Supplemental Figure 3D**).

Examination of PAD-associated module composition revealed the enrichment of bacteria with known functional significance. For example, the red module is enriched with known butyrate producers such as *Faecalibacterium*, *Faecalibacillus* and *Agathobaculum* genera (**Supplemental Figure 4**). Members of the modules that are negatively correlated with PAD severity such as cyan, red, pink and purple modules had negative correlations between kWithin and taxa significance (data not shown), which is defined by the correlation between taxa clr abundance and PAD severity. This indicates that the taxa directly contributing to PAD severity are peripherally located in the module, whereas central hubs maintain structural stability. This is further supported by the loss of Zdensity in these modules compared to the non-PAD network (**Supplemental Figure 3D**), suggesting the loss of peripheral members as the severity of PAD increases. The magenta module, which correlates positively with PAD severity and negatively with fecal butyrate levels, mostly consists of oral microbes such as *Granulicatella* spp, *S. oralis* clades, *Streptococcus*, and *Actinomyces* (**Supplemental Figure 4**). Most of the top network hubs are also module hubs in the turquoise and red modules (**Supplemental Table 2**), and they are also the main contributors driving variation in PC1 (**Figure 4C**).

Similarly, the module hubs in the top 25^th^ percentile of the purple (12/20 hubs), red (10/24 hubs) and turquoise (24/56 hubs) modules make up 46% of the top 100 contributors of PC1 variation (**Supplemental Table 2**). The observation that specific bacteria exhibit both high connectivity within co-occurrence networks and significant variation in abundance suggests that they act as key drivers of community shifts in PAD. Specifically, members of the turquoise module such as *Ruminococcus B sp 900544395*, *Blautia hansenii*, *Faecalimonas sp900546325*, *Clostridium AQ innocuum*, *Blautia sp001304935*, *Enterocloster clostridioformis* and *Clostridium AP scindens* represent top module hubs, top PC1 contributors, differentially abundant species in CLTI vs. claudication (**Figure 5A**) and fall within the top 4% of the network hubs, which further emphasizes their potential roles in disease progression.

### 8. Functional predictions

We identified 22 MetaCyc pathways including CENTFERM-PWY: pyruvate fermentation to butanoate that were reduced in the PAD compared to non-PAD groups (**Supplemental Figure 5A**). Similarly, the fatty acid biosynthetic pathways including SCFA such as CENTFERM-PWY (pyruvate fermentation to butanoate), PWY-6285 (superpathway of fatty acids biosynthesis), and PWY-6284 (superpathway of unsaturated fatty acids biosynthesis) were reduced in the CLTI compared to the claudication subgroups (**Figure 7A**), suggesting that the synthesis of SCFA including butyrate is disrupted in severe PAD. ^64^ Similarly, depletion in abundance of LIPA-CORESYN_PWY: lipid A-core biosynthesis (*E. coli* K-12) and LPSSYN-PWY: superpathway of lipopolysaccharide biosynthesis (**Figure 7A**) suggest reduced immune regulation in CLTI. In contrast, bile acid metabolism pathways such as PWY-8134 (bile acid 7β-dehydroxylation) and PWY-7754 (bile acid 7α-dehydroxylation) were increased in the CLTI subgroup (**Figure 7A** and **Figure 7B**). The main contributing species for the increased abundance of these secondary bile acid pathways in CLTI was *Clostridium scindens* (**Figure 7B** and **7C**). *Clostridium scindens* was grouped into the turquoise module in the PAD network that was associated with CLTI (**Supplement Figure 3C)** and negatively correlated with multiple clades of *F. prausnitzii* species in the red module (**Supplemental Figure 5B**) and *Roseburia hominis* and *Lachnospira eligens A* in the purple module, indicating *C. scindens’*s potential role in PAD progression by dysregulating secondary bile acid production and establishing negative associations with butyrate producers. Additionally, the node degree of *C. scindens* was smallest in the non-PAD network (node degree = 5) (**Figure 7D**) compared to the claudication (node degree = 16) (**Figure 7E**) and CLTI networks (node degree = 38) (**Figure 7F**), suggesting PAD-associated rewiring by increasing the negative correlation with butyrate producers.

**Figure 7.**
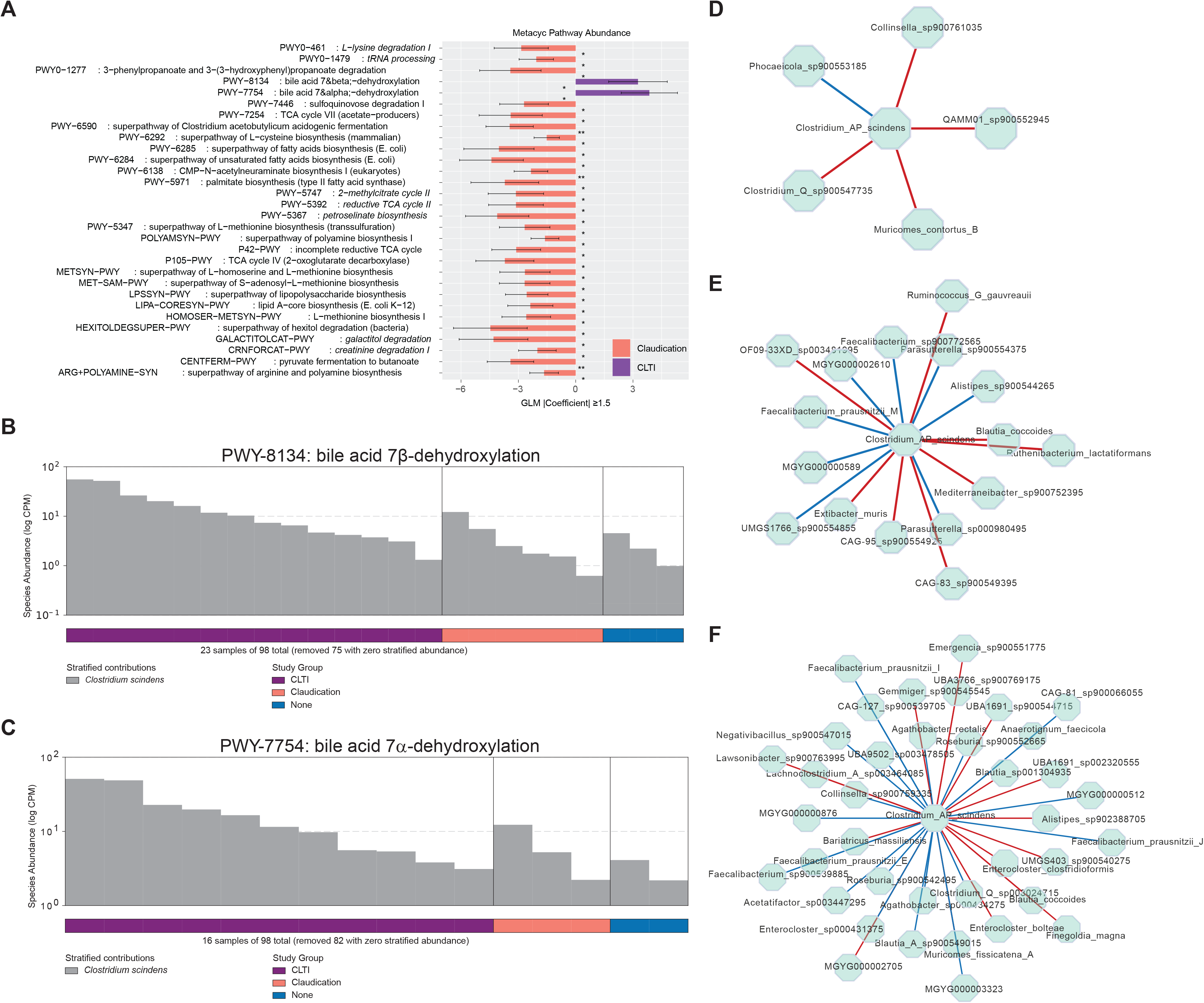
Pathway-level functional shift and network rewiring of corresponding taxa in the clinical groups. (A) Differentially abundant metabolic pathways between the claudication and CLTI subgroups. Pathways present in less than 10% of samples were excluded before analysis. *, p < 0.05. **, p < 0.01, Generalized linear model test. Species contribution to the differentially abundant secondary bile acid pathways, PWY-8134 (B) and PWY-7754 (C). Connections of Clostridium scindens to other species in the non-PAD (D), claudication (E), and CLTI (F) networks. Red and blue edges represent positive and negative correlations, respectively. Length of the edges were adjusted for better visualization of nodes and node labels.

## Discussion

In this cross-sectional study of adults with PAD and age- and sex-matched non-PAD controls, we used shotgun metagenomics and untargeted metabolomics to characterize PAD-associated gut dysbiosis as loss of microbial diversity, which becomes more pronounced as clinical severity of PAD progresses from claudication symptoms to chronic limb-threatening ischemia. We also identified patterns of differential abundance at the species level, network modules and hub species, and metabolic pathways associated with clinical severity of PAD.

Other cardiovascular diseases have been reported to be associated with gut dysbiosis, including coronary artery disease, stroke, and symptomatic carotid stenosis. A study using 16S rRNA amplicon sequencing showed decreased diversity and richness, enrichment of pathogenic bacteria such as *Escherichia-Shigella,* and depletion of butyrate producers such as *Roseburia* and *Eubacterium rectale* among people with coronary artery disease (CAD) compared to healthy controls. ^65^ A separate study identified correlations between specific gut bacterial taxa and coronary plaque phenotype and burden. ^66^ In a metagenomics and metabolomics study of people with stable CAD and diabetes mellitus, individuals with both conditions had increased levels of aromatic amino acids and carbohydrates, which correlated with a gut microbiome with enriched biosynthetic potential, while *Ruminococcus torques* was associated with cardiac survival. ^67^ Among patients with acute stroke, specific dysbiosis patterns across 18 genera were used to create a Stroke Dysbiosis Index (SDI) which was associated with stroke severity and outcome at hospital discharge. Fecal transplantation of mice from high-SDI donors was associated with more severe brain injury and inflammation following middle cerebral artery occlusion compared to low-SDI donors, suggesting a causal role of dysbiosis in stroke outcomes. ^68^ A separate study found that enrichment in Enterobacteriaceae was an independent risk factor for poor outcomes after acute stroke. ^69^ Intestinal ischemia caused by brain ischemia may result in gut dysbiosis, which could be treated by suppressing Enterobacteriaceae overgrowth in a mouse model of acute stroke. ^69^ In another study, the genus *Collinsella* was enriched in patients with symptomatic carotid stenosis, whereas *Roseburia* and *Eubacterium* were enriched in healthy controls. Patient gut metagenomes were also enriched in genes encoding peptidoglycan synthesis and depleted in phytoene dehydrogenase. ^42^

The unexpected lack of a notable shift in gut microbial community between non-PAD and PAD groups in our study may be due to the characteristics of our control group. Our control group was age- and sex-matched and included participants with cardiovascular risk factors such as hypertension, hyperlipidemia, obesity, smoking, and diabetes mellitus. Atherosclerosis is a disease of aging^70^ and is sexually dimorphic. ^71^ Age-related changes in gut microbiota composition, function, and diversity and in intestinal function and integrity are well-described, ^72-80^ as are sex-based differences in the gut microbiome across multiple species. ^81-87^ By including participants with cardiovascular risk factors in the control cohort rather than using “healthy” controls as in other studies, ^88 53^ we were able to examine dysbiosis associated specifically with PAD and avoid confounding by these risk factors, which are also known to be associated with dysbiosis. ^89-104^ In fact, in our cohort, significant differences between the non-PAD and PAD groups were not seen following adjustment for age, sex, and comorbidities. Interestingly, the overlap between many of the strongly positive species loadings in PC1 and the differentially abundant species between the PAD clinical subgroups suggest that PC1 is driven by the composition differences between the claudication and CLTI subgroups more than those between the non-PAD and PAD groups. Higher PC1 scores in the claudication subgroup compared to the non-PAD group suggest a transient increase in the relative abundance of beneficial bacteria for gut homeostasis followed by their depletion in CLTI. Similarly, relative enrichment of butyrate producers in claudication suggests a compensation in beneficial metabolic functions before advancing into a more dysbiotic state associated with CLTI.

Butyrate production is widespread among diverse colonic phyla, ^105^ but are mainly in Firmicutes and Bacteroides. ^62^ The most prominent butyrogenic species are *F. prausnitzii*, *Butyrivibrio crossotus* and *Roseburia intestinalis*. ^62^ Butyrate is known to be a histone deacetylase inhibitor, ^106^ agonist of peroxisome proliferator-activated receptors (PPARs) and free fatty acid receptors, ^107-112^ and activator of AMP kinase, which upregulates PPARγ coactivator, PPARα and PPARγ. ^109,113^ We have previously demonstrated that microbe-associated butyrate reduces neointimal hyperplasia after arterial surgery in a mouse model. ^114^ Others have found an inverse association between butyrate levels, butyrate producer abundance, and other cardiovascular diseases. ^42,62,115-124^ In concordance with the above studies, we found that butyrate producers are associated with clinically less severe PAD. Furthermore, other butyrate producers such as *Butyricimonas virosa* and *Intestinimonas butyriciproducens* are less abundant in both PAD subgroups compared to the non-PAD group. The abundance of pathways related to the fermentation of pyruvate and acetyl-CoA to butanoate are also differentially reduced in CLTI compared to non-PAD. We did not find a statistically significant difference in the abundance of butyrogenic pathways between claudicants and the non-PAD group, suggesting that both butyrate production pathways become less abundant only in CLTI, the most clinically advanced form of PAD.

Bile acid metabolism is dependent on gut microbes. ^125-131^ Bile acids are key contributing factors to lipid homeostasis and glucose metabolism^132^ and have been shown to cause intestinal inflammation, colon cancer^133^ and increase the risk of cardiovascular diseases. ^134^ In the current study, we found that bile acid 7α- and 7β-dehydroxylation pathways in the CLTI subgroup are upregulated compared to the claudication subgroup, and the abundance is mainly driven by *Clostridium scindens*, which is regarded as a keystone species in the human colon for major biotransformations of bile acids and other steroids that regulate the structure of the gut microbiome and host–microbe interactions. ^135^ *C. scindens* was also differentially abundant in the CLTI compared to the claudication subgroup and has negative co-occurrence with butyrate producers such as *F. prausnitzii* clades, implying that *C. scindens* contributes to PAD severity by outcompeting or disrupting the balance of beneficial microbes. Secondary bile acid, deoxycholic acid (DCA), produced by *C. scindens* has been shown to inhibit the growth of *F. prausnitzii* ^136^ and to impair glucose metabolism and worsen lipid profiles in experimental mice^137^ Colonization of germ-free mice with *F. prausnitzii-containing communities* lowered the intestinal concentration of DCA primarily by reducing the expression of bile acid-CoA transferase/hydrolase and BNADH-dependent oxidoreductase genes^136^ that are involved in the reductive arm in the 7α-dehydroxylation pathway. ^135^ This can potentially create gut microbiome homeostasis by maintaining the optimum DCA level and increasing bioactive intermediate products.

The gut microbiome is a complex microecosystem of constantly interacting communities. Previous studies have found altered microbial interaction patterns in atherosclerotic cardiovascular disease. ^8^ Our results also indicate a dynamic reorganization of the gut microbiome interaction pattern in PAD. Community interconnectedness increases in claudication, an early disease state (**Figure 6A**), suggesting an adaptive response to environmental stress, which is followed by the loss of connectivity, indicating community disintegration in CLTI, or severe PAD. Our findings complement non-linear stress-response-collapse dynamics of microbial communities^138 139^ similar to other complex ecosystems. ^140^ However, rewiring of connectivity happened simultaneously with progressive fragmentation of clusters (**Figure 6A**). In other studies, fragmentation and rewiring of connectivity patterns are linked to greater susceptibility of inflammation and disease such as infant respiratory tract infection, erosive gastrointestinal inflammation and antibiotic-induced inflammatory response. ^141 142 62^ In our study, the affected clusters are mostly the SCFA producers that are negatively associated with PAD severity (**Supplemental Figure 3C**). These groups of beneficial taxa became more densely interconnected within their clusters, possibly to limit global interaction and avoid extinction, as suggested previously. ^138^ We also observed that the difference in community richness between non-PAD and PAD clinical stages may be sex-specific (**Supplement Figure 1**). This indicates a different mechanism of community destabilization between the sexes. For instance, in males, an increase in richness may precede the destabilization process. ^138^

To our knowledge, this is the most comprehensive evaluation of the gut microbiome and metabolites in patients with PAD and its clinical subgroups to date. Prior studies have focused on cardiovascular death among adults with PAD, ^11^ used 16S rRNA amplicon sequencing in a smaller cohort, ^143^ or did not include metabolomics. ^12^ The finding of gut microbial community features associated with various stages of PAD can be used to investigate potential microbiome biomarkers for early diagnosis or intervention or microbe-related preventative strategies such as lifestyle and dietary changes or pre- and probiotic supplements. Additional strengths include the multi-omics approach, which addresses a substantial knowledge gap in the functional characterization of gut microbiota in PAD and justifies further focused investigation using a larger cohort. Finally, the use of an age- and sex-matched non-PAD cohort and adjustment for atherosclerosis risk factors rigorously addresses confounding variables.

There are several limitations to this study. First, the overall cohort size is modest and the smaller size of the non-PAD control group compared to the PAD group potentially obscures community level differences driven by rare species. Second, metabolomics was performed using only a subset of randomly selected samples and thus does not represent a full correlation with metagenomics dataset. Third, we utilized metabolomics using NMR, which is quantitative, reproducible, non-destructive and less costly compared to mass spectrometry but has lower sensitivity and detects fewer metabolites. ^144^ However, our findings can be used as a starting point for future targeted analyses or to justify future analyses using mass spectrometry that may reveal other metabolites associated with PAD. Fourth, we did not account for dietary habits in our study. Among community-dwelling middle-aged adults, meat consumption is associated with increased incident PAD over 20 years of follow-up, while coffee consumption and alcohol are inversely associated with incident PAD. ^145^ Furthermore, daily consumption of greater quantities of fruit and vegetables is associated lower odds of PAD. ^146^ Since prior work has demonstrated shifts in microbiome diversity, composition, and microbial metabolites based on animal-based or plant-based diets, ^147-151^ it will be important for future characterizations of PAD-associated dysbiosis to include dietary information.

In conclusion, we characterize dysbiosis in PAD as loss of diversity and increased homogeneity, while CLTI correlates with reduced SCFA producers and rewiring of the gut microbiota co-occurrence pattern. These specific changes to the gut community suggest that microbe-mediated therapy could potentially modulate progression of PAD or that PAD prognosis can be personalized based on microbiome-features. The sensitivity of gut microbiome networks to abundance fluctuations offers promising avenues for targeted ecosystem interventions in PAD. Future studies utilizing a larger and clinically well-annotated cohort will validate our observations, while experimental models will uncover the biological mechanisms and pathways for the role of gut microbes in PAD pathogenesis and progression.

## Declarations

### Ethics approval and consent to participate

This study was conducted in accordance with the Declaration of Helsinki. All participants provided written informed consent. This study was approved by the Northwestern University Institutional Review Board (Protocols #STU00213201 and #STU00216722).

### Consent for publication

Not applicable (no data from any individual person)

### Availability of data and materials

The datasets generated and/or analyzed during the current study are in a publicly available repository (NCBI Sequence Read Archive) under BioProject accession number PRJNA1457558.

### Authors’ contributions statement

All authors made substantial contributions to the conception, design of the work, acquisition, analysis, or interpretation of data; assisted in the drafting or revision of the manuscript; approved the submitted version, and have agreed both to be personally accountable for the author’s own contributions and to ensure that questions related to the accuracy or integrity of any part of the work, even ones in which the author was not personally involved, are appropriately investigated, resolved, and the resolution documented in the literature.

## Acknowledgements

Not applicable

## List of Abbreviations and Acronyms

BH: Benjamini-Hochberg
BMI: body mass index
CLTI: chronic limb-threatening ischemia CPM copies per million
kME: module membership
kOut: Iintermodular connectivity kWithin intramodular connectivity ME module eigengenes
NMR: nuclear magnetic resonance
PAD: peripheral artery disease
PC: principal component TMSP trimethylsilylpropanoic acid
WGCNA: weighted gene co-expression network analysis

