## Supplement figures and legends for "Shotgun metagenomic and untargeted metabolomic analysis of gut microbiome associated with peripheral artery disease"

1. Supplemental Figure 1-5
2. Supplemental Table 1
3. Supplemental Table 2

**Supplemental Figure 1. Sexual dimorphism in alpha diversity in the clinical groups.** (A) Comparison of alpha diversity indices between the non-PAD and PAD groups stratified by sex. (B) Comparison of alpha diversity indices between the non-PAD, claudication and CLTI subgroups stratified by sex. *, $p< 0.05$, Wilcoxon rank-sum test

**Supplemental Figure 2. Untargeted metabolomics of fecal samples from the non-PAD and PAD groups.** (A) Box and whiskers plot of log-transformed metabolite concentrations in the non-PAD and PAD groups. Metabolites are grouped into biochemical categories for ease of visualization. The boxes represent the 25% and 75% percentiles. Upper and lower limit values are indicated by the horizontal lines. Outliers are not shown. Median values are indicated by vertical lines within each box. (B) Box and whiskers plot of log-transformed short chain fatty acid concentrations stratified by group and presenting symptoms, including outlier values. *, $p < 0.05$, Wilcoxon rank-sum test. UDP, Uridine diphosphate. 3-HPA, 3-Hydroxyphenylacetate.

**Supplemental Figure 3. Network module relationships with each other and with sample traits.** Circose plots showing positive (A) and negative (B) correlations among modules in the PAD network. Each color represents a different module. Ribbon thickness represents the average strength of the connections and opacity represents average strength of the connection. (C) Spearman correlation of module eigengenes (vertical axis) with sample traits. Vertical axis module dendrogram represents hierarchical clustering of module eigengenes using the average linkage method. Traits are listed on the horizontal axis. The “CLTI” trait refers to correlation with claudication (0) and CLTI (1). The “Symptom” trait refers to the presenting symptom of PAD (claudication, 0; rest pain 1; tissue loss 2). (D) Preservation of the PAD network module density (left panel) and connectivity (right panel) within the non-PAD network. Red and blue dashed lines represent lower (Z=2) and upper (Z=10) threshold values for moderate z-statistics.

**Supplemental 4.** Enriched genera in each module with respect to all species in the PAD network. The enrichment score was calculated based on Fisher’s Z test and is represented by the length of the horizontal bars. Color gradients represent $\log_{10}$ q-values that were BH-adjusted across modules. Bubble size represents proportion of a specific genus within a module compared to its abundance across the entire network.

**Supplemental Figure 5.** (A) Differences in MetaCyc pathway abundances between the non-PAD and PAD group. Differentially more abundant pathways were found only in the non-PAD group. *, $p < 0.05$. **, $p < 0.01$, Generalized linear model test. (B) Connections of *Clostridium AP scindens* with members of other modules. Circles represent species. Circle size indicates the taxa significance. Red and blue lines represent positive and negative correlations, respectively. Length of the lines were adjusted for better visualization of nodes and node labels.

**Supplemental Table 1. Sample Metadata and Sequence Processing Summary.**

**Supplemental Table 2. Module statistics for PAD network.**

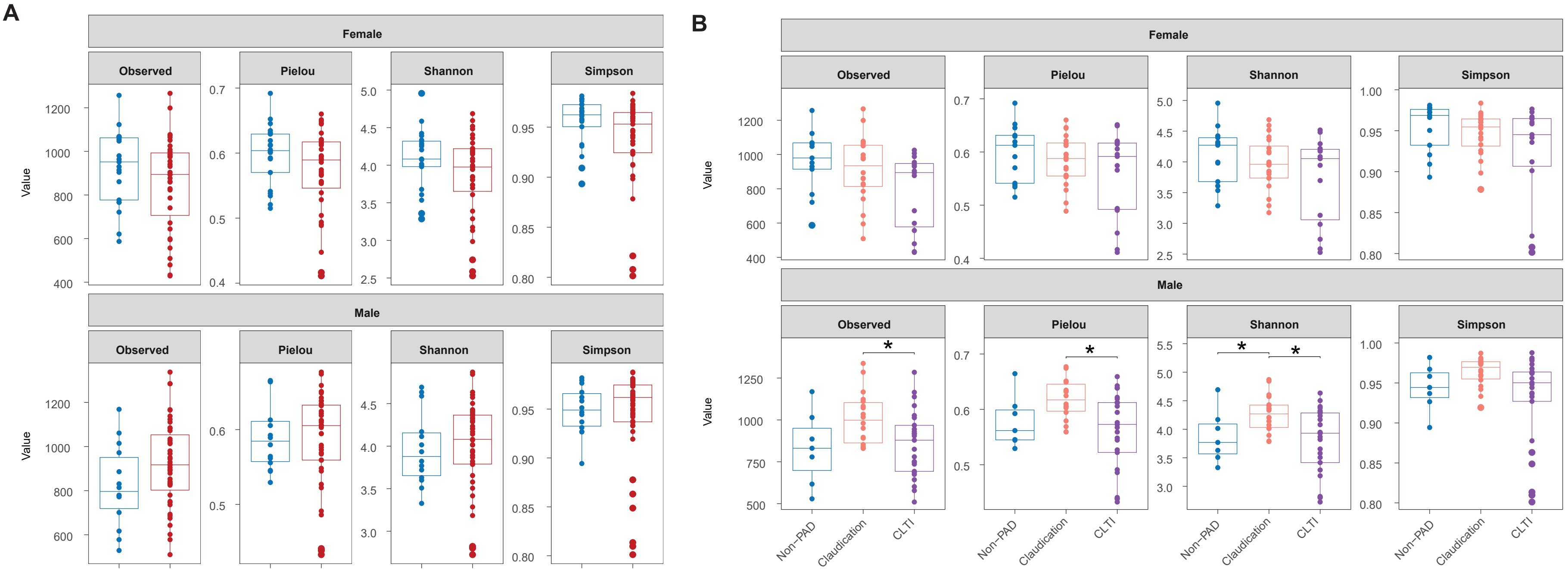
A
Female
Observed
Pielou
Shannon
Simpson
Male
Value
Non−PAD
PAD
B
Claudication
CLTI
*


Supp Fig 1

A

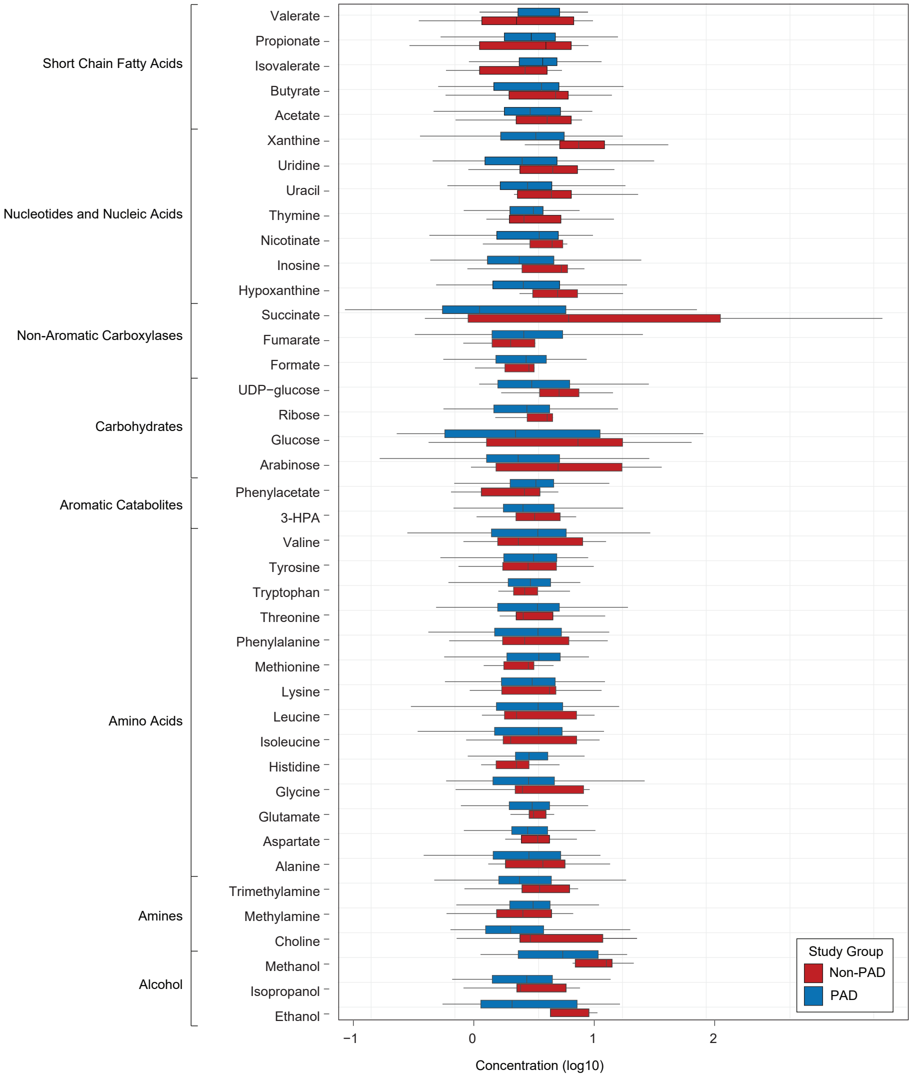

Short Chain Fatty Acids
Nucleotides and Nucleic Acids
Non-Aromatic Carboxylases
Carbohydrates
Aromatic Catabolites
Amino Acids
Amines
Alcohol
Valerate
Propionate
Isovalerate
Butyrate
Acetate
Xanthine
Uridine
Uracil
Thymine
Nicotinate
Inosine
Hypoxanthine
Succinate
Fumarate
Formate
UDP−glucose
Ribose
Glucose
Arabinose
Phenylacetate
3-HPA
Valine
Tyrosine
Tryptophan
Threonine
Phenylalanine
Methionine
Lysine
Leucine
Isoleucine
Histidine
Glycine
Glutamate
Aspartate
Alanine
Trimethylamine
Methylamine
Choline
Methanol
Isopropanol
Ethanol
−1
0
1
2
Concentration (log10)
Study Group
Non-PAD
PAD


B

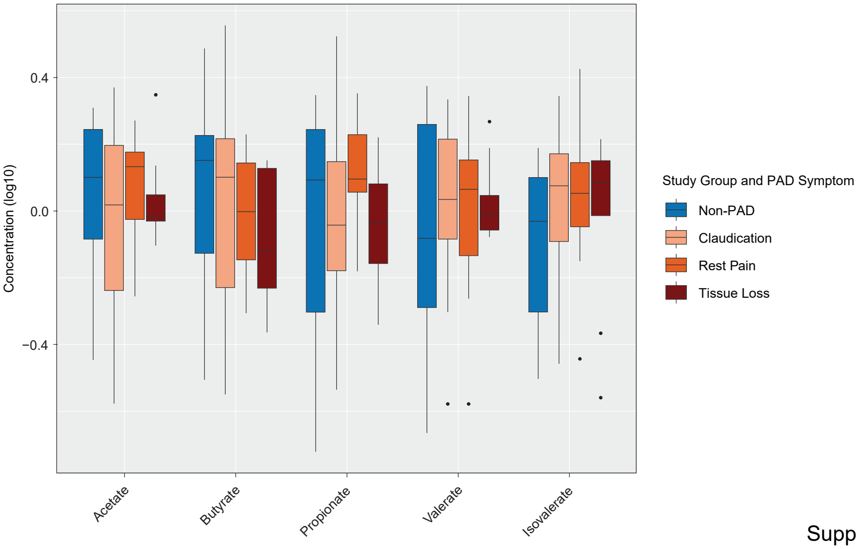

0.4
0.0
−0.4
Concentration (log10)
Acetate
Butyrate
Propionate
Valerate
Isovalerate
Study Group and PAD Symptom
Non-PAD
Claudication
Rest Pain
Tissue Loss


Supp Fig 2

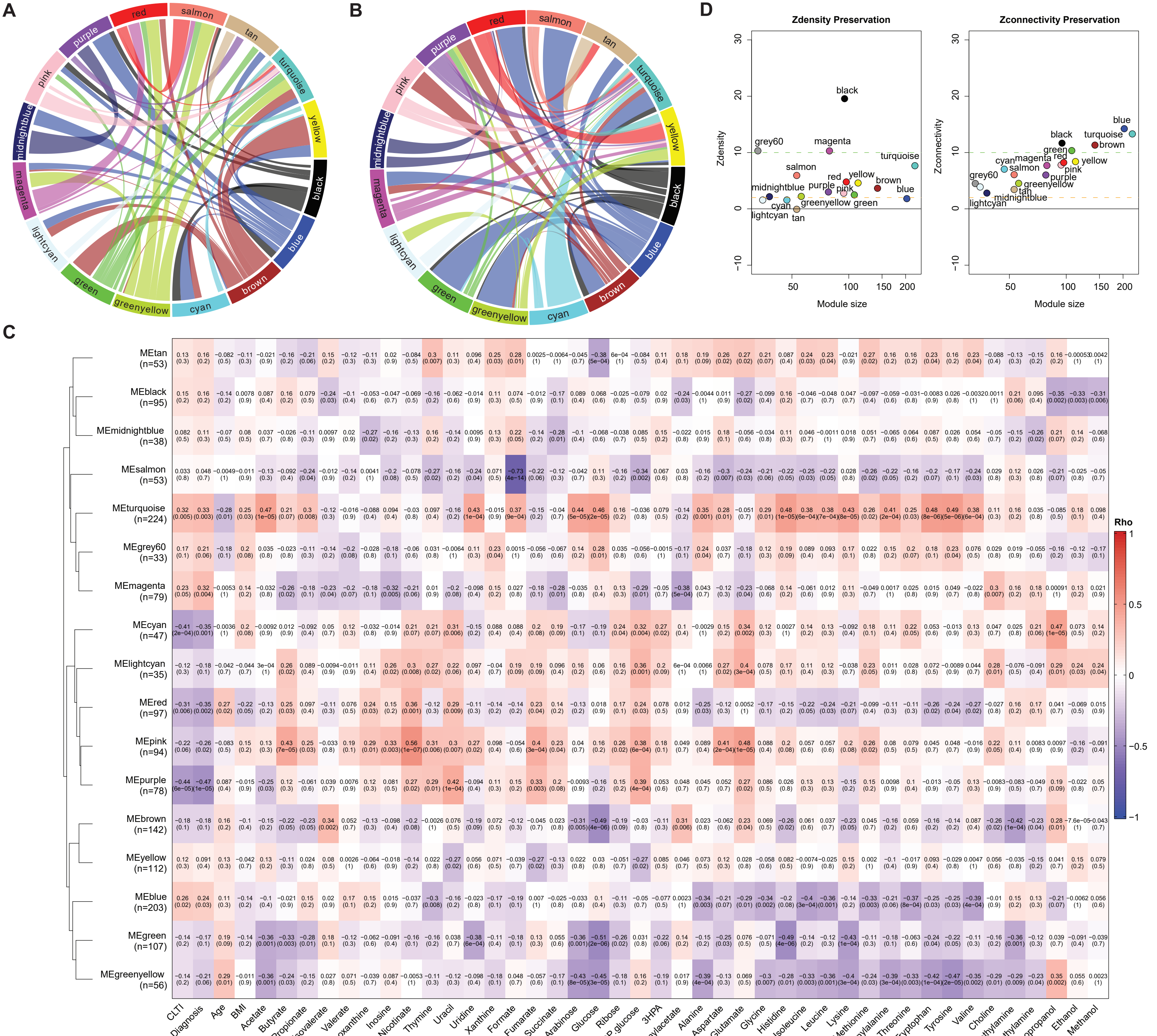
A
B
C
D
red
salmon
tan
turquoise
yellow
black
blue
brown
cyan
greenyellow
green
lightcyan
magenta
midnightblue
pink
purple
Zdensity Preservation
Zconnectivity Preservation
Zdensity
Zconnectivity
Module size
grey60
Rho
MEtan (n=53)
MEblack (n=95)
MEmidnightblue (n=38)
MEsalmon (n=53)
MEturquoise (n=224)
MEgrey60 (n=33)
MEmagenta (n=79)
MEcyan (n=47)
MElightcyan (n=35)
MEred (n=97)
MEpink (n=94)
MEpurple (n=78)
MEbrown (n=142)
MEyellow (n=112)
MEblue (n=203)
MEgreen (n=107)
MEgreenyellow (n=56)
CLTI
Diagnosis
Age
BMI
Acetate
Butyrate
Propionate
Isovalerate
Valerate
Hypoxanthine
Inosine
Nicotinate
Thymine
Uracil
Uridine
Xanthine
Formate
Fumarate
Succinate
Arabinose
Glucose
Ribose
UDP.glucose
3HPA
Phenylacetate
Alanine
Aspartate
Glutamate
Glycine
Histidine
Isoleucine
Leucine
Lysine
Methionine
Phenylalanine
Threonine
Tryptophan
Tyrosine
Valine
Choline
Methylamine
Trimethylamine
Isopropanol
Ethanol
Methanol

Supp Fig 3

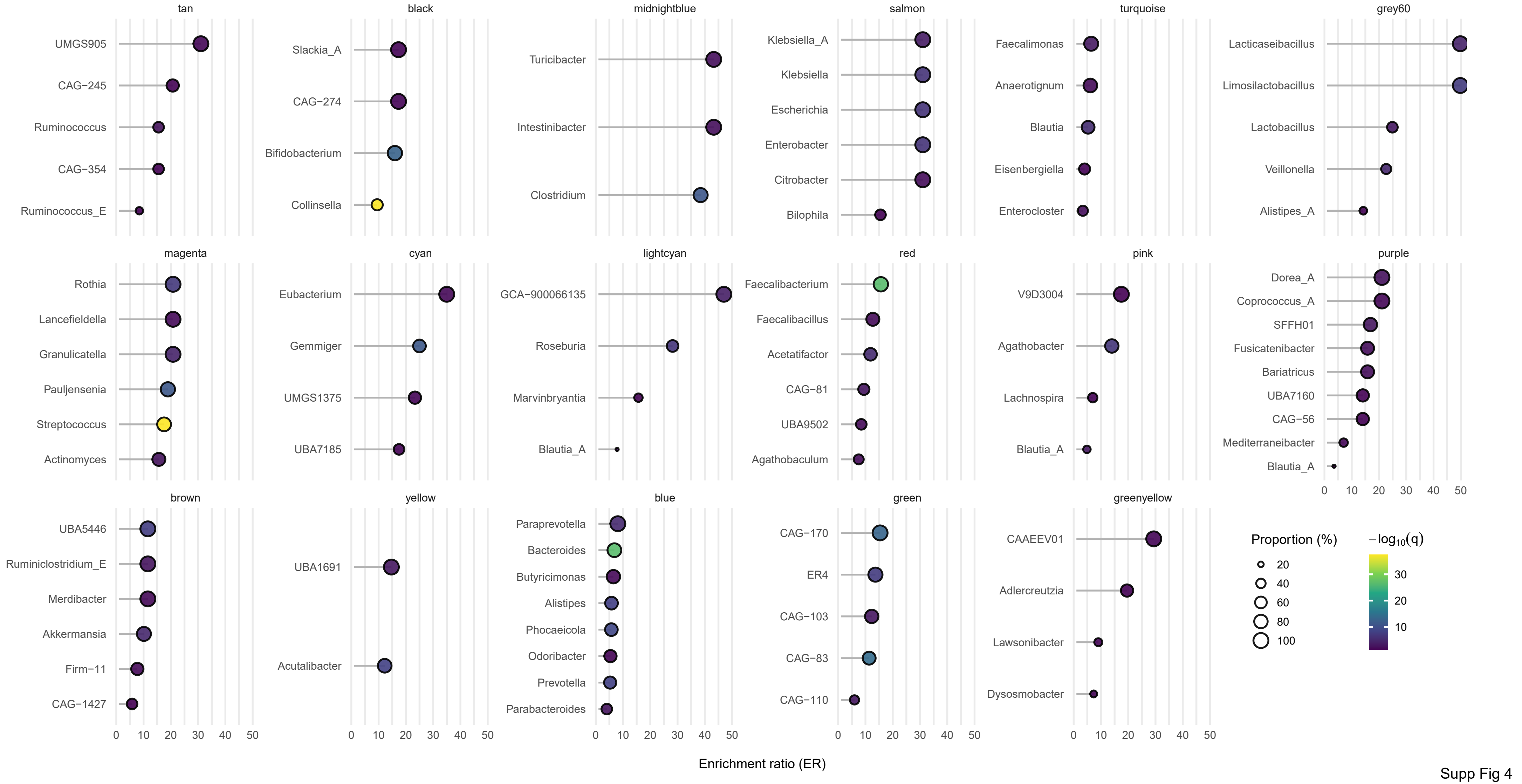
tan
UMGS905
CAG−245
Ruminococcus
CAG−354
Ruminococcus_E
black
Slackia_A
CAG−274
Bifidobacterium
Collinsella
midnightblue
Turicibacter
Intestinibacter
Clostridium
salmon
Klebsiella_A
Klebsiella
Escherichia
Enterobacter
Citrobacter
Bilophila
turquoise
Faecalimonas
Anaerotignum
Blautia
Eisenbergiella
Enterocloster
grey60
Lacticaseibacillus
Limosilactobacillus
Lactobacillus
Veillonella
Alistipes_A
magenta
Rothia
Lancefieldella
Granulicatella
Pauljensenia
Streptococcus
Actinomyces
cyan
Eubacterium
Gemmiger
UMGS1375
UBA7185
lightcyan
GCA−900066135
Roseburia
Marvinbryantia
Blautia_A
red
Faecalibacterium
Faecalibacillus
Acetatifactor
CAG−81
UBA9502
Agathobaculum
pink
V9D3004
Agathobacter
Lachnospira
Blautia_A
purple
Dorea_A
Coprococcus_A
SFFH01
Fusicatenibacter
Bariatricus
UBA7160
CAG−56
Mediterraneibacter
Blautia_A
brown
UBA5446
Ruminiclostridium_E
Merdibacter
Akkermansia
Firm−11
CAG−1427
yellow
UBA1691
Acutalibacter
blue
Paraprevotella
Bacteroides
Butyricimonas
Alistipes
Phocaeicola
Odoribacter
Prevotella
Parabacteroides
green
CAG−170
ER4
CAG−103
CAG−83
CAG−110
greenyellow
CAAEEV01
Adlercreutzia
Lawsonibacter
Dysosmobacter
0 10 20 30 40 50
Proportion (%)
20
40
60
80
100
$-\log_{10}(q)$
30
20
10
Enrichment ratio (ER)

Supp Fig 4

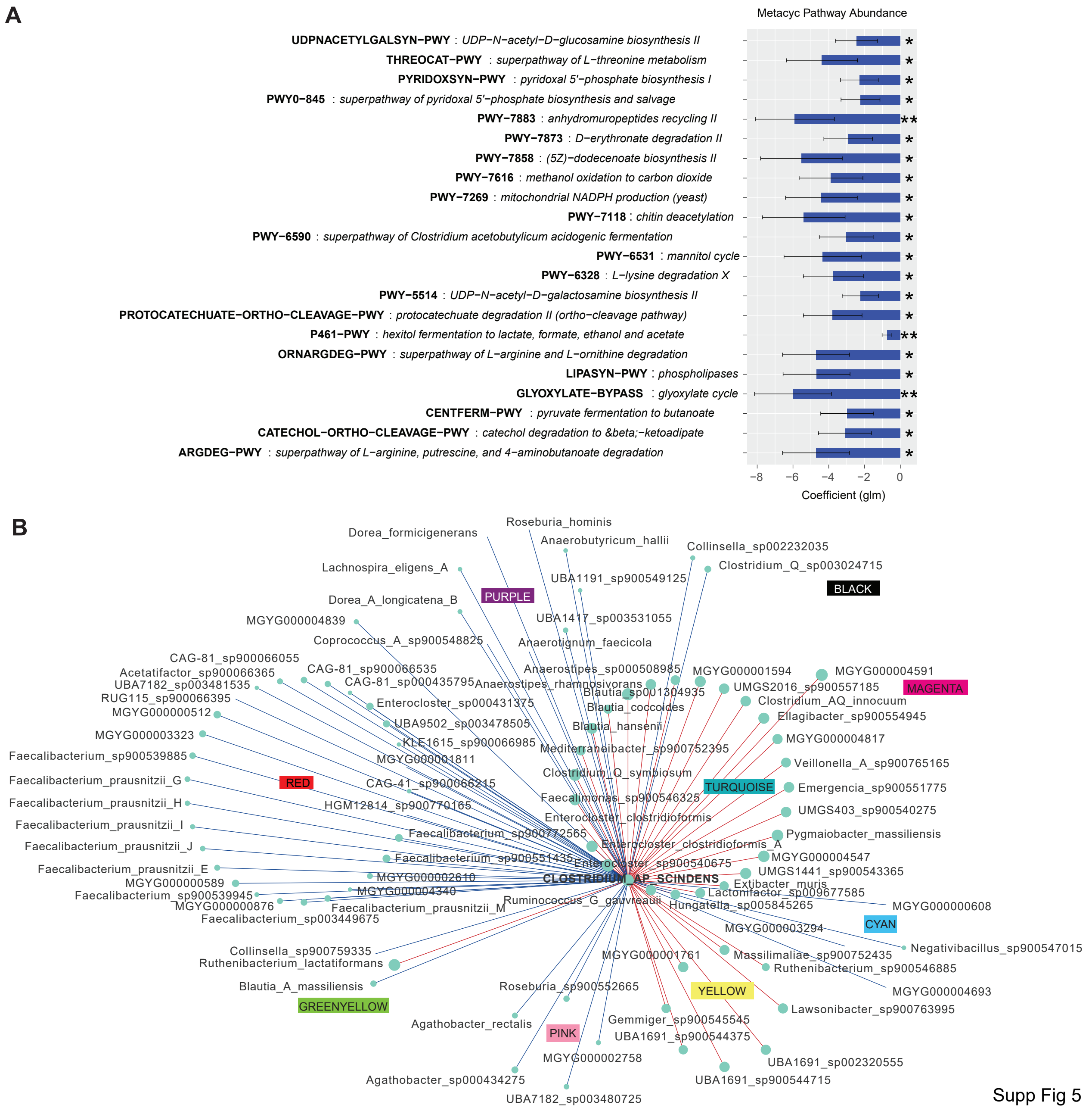
A
Metacyc Pathway Abundance
UDPNACETYLGALSYN−PWY : UDP−N−acetyl−D−glucosamine biosynthesis II
THREOCAT−PWY : superpathway of L−threonine metabolism
PYRIDOXSYN−PWY : pyridoxal 5'−phosphate biosynthesis I
PWY0−845 : superpathway of pyridoxal 5'−phosphate biosynthesis and salvage
PWY−7883 : anhydromuropeptides recycling II
PWY−7873 : D−erythronate degradation II
PWY−7858 : (5Z)−dodecenoate biosynthesis II
PWY−7616 : methanol oxidation to carbon dioxide
PWY−7269 : mitochondrial NADPH production (yeast)
PWY−7118 : chitin deacetylation
PWY−6590 : superpathway of Clostridium acetobutylicum acidogenic fermentation
PWY−6531 : mannitol cycle
PWY−6328 : L−lysine degradation X
PWY−5514 : UDP−N−acetyl−D−galactosamine biosynthesis II
PROTOCATECHUATE−ORTHO−CLEAVAGE−PWY : protocatechuate degradation II (ortho−cleavage pathway)
P461−PWY : hexitol fermentation to lactate, formate, ethanol and acetate
ORNARGDEG−PWY : superpathway of L−arginine and L−ornithine degradation
LIPASYN−PWY : phospholipases
GLYOXYLATE−BYPASS : glyoxylate cycle
CENTFERM−PWY : pyruvate fermentation to butanoate
CATECHOL−ORTHO−CLEAVAGE−PWY : catechol degradation to β−ketoadipate
ARGDEG−PWY : superpathway of L−arginine, putrescine, and 4−aminobutanoate degradation
−8 −6 −4 −2 0
Coefficient (glm)
B
PURPLE
BLACK
MAGENTA
RED
TURQUOISE
CYAN
GREENYELLOW
PINK
YELLOW
CLOSTRIDIUM_AP_SCINDENS
Roseburia_hominis
Dorea_formicigenerans
Anaerobutyricum_hallii
Collinsella_sp002232035
Clostridium_Q_sp003024715
Lachnospira_eligens_A
UBA1191_sp900549125
Dorea_A_longicatena_B
UBA1417_sp003531055
MGYG000004839
Coprococcus_A_sp900548825
Anaerotignum_faecicola
CAG-81_sp900066055
CAG-81_sp900066535
Anaerostipes_sp000508985
MGYG000001594
MGYG000004591
Acetatifactor_sp900066365
CAG-81_sp000435795
Anaerostipes_rhamnosivorans
UBA7182_sp003481535
Blautia_sp001304935
UMGS2016_sp900557185
RUG115_sp900066395
Enterocloster_sp000431375
Clostridium_AQ_innocuum
MGYG000000512
Blautia_coccoides
Ellagibacter_sp900554945
UBA9502_sp003478505
Blautia_hansenii
MGYG000003323
KLE1615_sp900066985
MGYG000004817
Faecalibacterium_sp900539885
MGYG000001811
Mediterraneibacter_sp900752395
Veillonella_A_sp900765165
Clostridium_Q_symbiosum
Faecalibacterium_prausnitzii_G
CAG-41_sp900066215
Emergencia_sp900551775
Faecalimonas_sp900546325
Faecalibacterium_prausnitzii_H
HGM12814_sp900770165
UMGS403_sp900540275
Enterocloster_clostridioformis
Faecalibacterium_prausnitzii_I
Faecalibacterium_sp900772565
Pygmaiobacter_massiliensis
Faecalibacterium_prausnitzii_J
Enterocloster_clostridioformis_A
Faecalibacterium_sp900551435
MGYG000004547
Enterocloster_sp900540675
Faecalibacterium_prausnitzii_E
UMGS1441_sp900543365
MGYG000002610
MGYG000000589
Extibacter_muris
MGYG000004340
Lactonifactor_sp009677585
Faecalibacterium_sp900539945
Ruminococcus_G_gauvreauii
MGYG000000608
MGYG000000876
Faecalibacterium_prausnitzii_M
Hungatella_sp005845265
Faecalibacterium_sp003449675
MGYG000003294
Negativibacillus_sp900547015
Collinsella_sp900759335
MGYG000001761
Massilimaliae_sp900752435
Ruthenibacterium_lactatiformans
Ruthenibacterium_sp900546885
Blautia_A_massiliensis
Roseburia_sp900552665
MGYG000004693
Lawsonibacter_sp900763995
Agathobacter_rectalis
Gemmiger_sp900545545
UBA1691_sp900544375
MGYG000002758
UBA1691_sp002320555
Agathobacter_sp000434275
UBA1691_sp900544715
UBA7182_sp003480725

Supp Fig 5
